# Secondary nucleation of α-Synuclein drives Mitochondria dysfunctions and Lewy body formation in Parkinson’s Disease

**DOI:** 10.1101/2025.09.17.676873

**Authors:** Emily E. Prescott, Jiapeng Wei, Emma F. Garland, Agnieszka Urbanek, Francesco Capriglia, Willem H. Molenkamp, David P Rhodes, Louise Heywood, Aurelie Schwartzentruber, Elezabeth Stephen, Rachel Hughes, Oliver Bandmann, Mark O. Collins, Sara Linse, Axel Abelein, Jan Johansson, Michele Vendruscolo, J Robin Highly, Tuomas P.J. Knowles, Heather Mortiboys, Suman De

**Author notes:** Corresponding author (T.P.J. K.) (H.M.) (S.D.).

## Abstract

The seeding of α-Synuclein (αSyn) is a key driver of Lewy pathology propagation in Parkinson’s disease (PD) and forms the basis for recent diagnostic advances. However, it remains unclear how the structural and biochemical features of αSyn seeds dictate their propagation efficiency, capacity to induce Lewy body formation, and resulting cellular toxicity. Using genetic and idiopathic PD cell models, we map the pathogenic cascade beginning with the seed-driven conversion of endogenous αSyn, followed by impaired degradation, mitochondrial dysfunction, and ultimately Lewy body formation. By coupling kinetic modelling of aggregation with functional readouts, we identify secondary nucleation as the predominant mechanism generating toxic αSyn aggregation intermediates, identifying the critical process that links seeding to pathology. Extending this framework to PD brain, we quantitatively correlate seeding capacity with the spatiotemporal spread and severity of Lewy pathology, revealing a mechanistic connection between αSyn aggregation dynamics and disease progression at molecular, cellular, and anatomical levels. By unifying molecular mechanism with clinicopathological progression, our work identifies catalytic αSyn fibrillar seeds as tractable targets for both disease-modifying therapy and biomarker development in PD.

**Graphical abstract:** 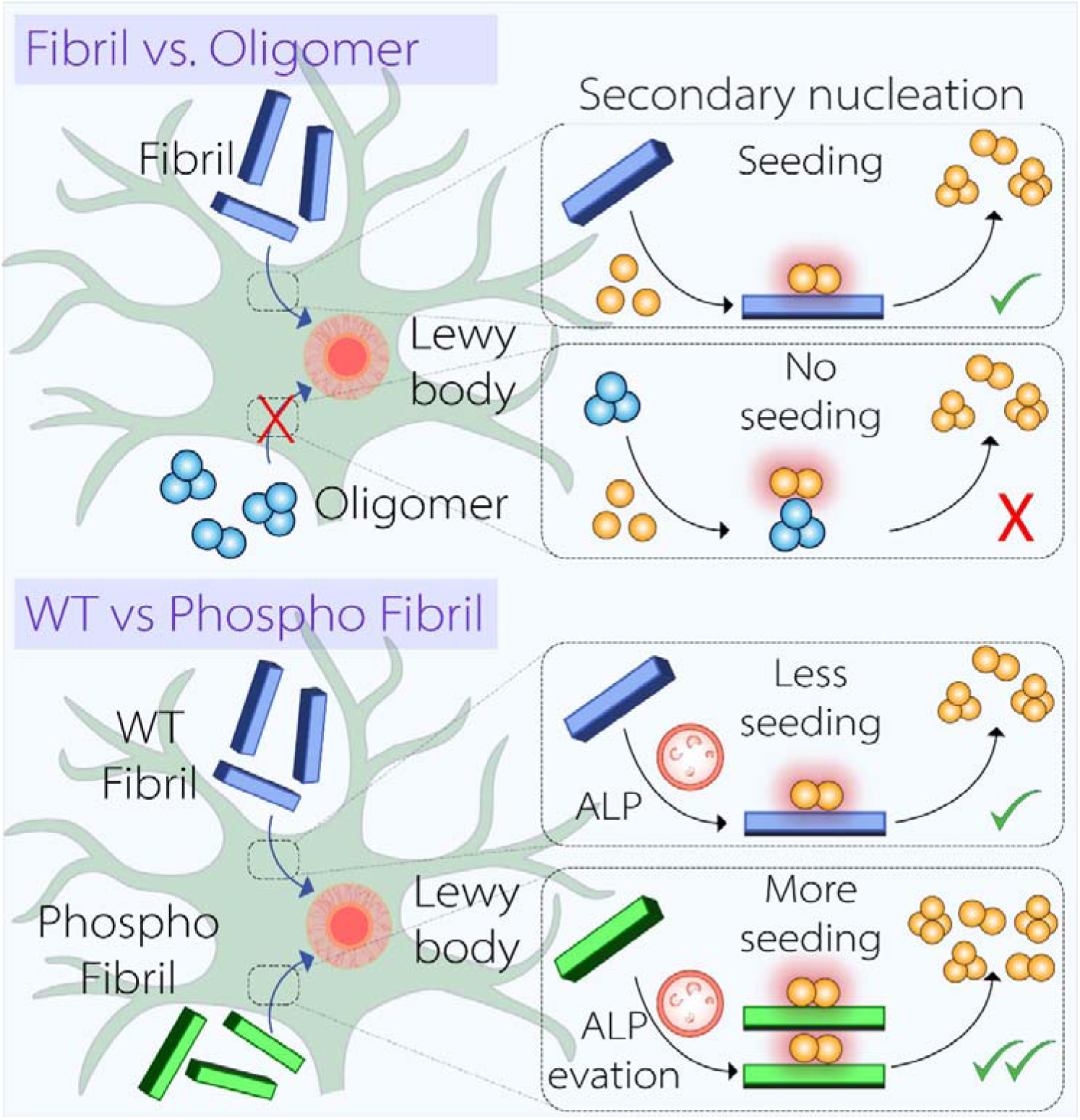

**Highlights:**

- αSyn fibril-oligomer interplay drives mitochondrial abnormalities and Lewy pathology
- Fibrillar αSyn catalyse toxic aggregate formations via secondary nucleation
- Phosphorylated αSyn evades lysosomal clearance and drives enhanced dysfunctions
- Seeding capacity of αSyn predicts Lewy pathology burden and disease progression

## Introduction

Parkinson’s disease (PD) is a progressive neurodegenerative disorder marked by debilitating motor symptoms such as bradykinesia, rigidity, and tremor^1,2^. While most cases are sporadic, approximately 5-10% are inherited genetic forms of the disease^3^. Among these, multiplications of the *SNCA* gene, which encodes α-synuclein (αSyn), result in increased αSyn expression and are associated with early-onset, rapidly progressing PD, establishing αSyn as a central player in disease pathogenesis^4–6^. In contrast, mutations in *PRKN*, which encodes the E3 ubiquitin ligase Parkin, typically cause autosomal recessive parkinsonism that often lacks classical αSyn pathology^7,8^. These two genetic forms, one with pronounced αSyn dysfunctions and the other largely devoid of it, reflect opposite ends of the pathology spectrum and raise fundamental questions about the causative role of αSyn in PD. Meanwhile, the majority of PD cases are idiopathic, with no defined genetic origin, and show high variability in clinical presentation, age of onset, and rate of progression^3,5,6^.

Despite the heterogeneity, almost all cases of PD converge on common pathogenic pathways: progressive degeneration of dopaminergic neurons in the Substantia Nigra, widespread dysfunction of subcellular compartments, particularly mitochondria and lysosomes and accumulation of αSyn-rich Lewy bodies (LBs) and Lewy neurites^5,9–11^. LBs are complex, multicomponent inclusions thought to form through a multistep process in which physiological αSyn misfolds, adopts β-sheet conformations, and assembles into higher-order species such as oligomers and fibrils, often accompanied by post-translational modifications^10,11^. These pathological assemblies disrupt key organelle functions, driving neuronal dysfunction and death^9,10,12^.

Yet, the molecular identity of αSyn species that drive LB formation and cause organelle dysfunction remains incompletely defined. Although fibrillar αSyn has long been recognised as the major constituent of LBs^11,13^, recent studies indicate that non-fibrillar forms may predominate in some cases^12^. Adding further complexity, post-translational modifications, particularly phosphorylation at serine 129 (pS129), present in up to 90% of LB-associated αSyn^14,15^. Although reducing pS129 levels has been shown to improve motor function in vivo^16^, some recent studies propose that pS129 may have physiological functions at synapses^17,18^, raising the question of whether pS129 promotes pathology or merely marks it.

A defining feature of PD is the spatiotemporal spread of Lewy pathology through anatomically connected brain regions, that closely mirrors clinical progression^19,20^. In this propagation misfolded αSyn seeds convert physiological αSyn into pathological aggregates forms^21,22^. These newly formed aggregates not only disrupt organelles and contribute to LB formation but also propagate to neighbouring neurons, where they trigger further αSyn aggregation and sustain the spread of pathology^23–25^. This concept underpins the development of αSyn seed amplification assays, which enable ante-mortem diagnosis with high sensitivity^26^. Moreover, faster amplification rates in this assay have been associated with increased disease risk^27^, and severity^28^, suggesting that αSyn seeding capacity may be a key determinant of disease trajectory.

However, it remains unclear whether heightened αSyn seeding activity is a mechanistic driver of disease or merely a diagnostic biomarker. Does elevated seeding capacity actively drive LB formation and disease progression, or does it simply reflect disease presence independent of LB burden? Is this process dependent on total αSyn levels, as observed in *SNCA* multiplication cases where gene dosage influences pathology, or can it arise independently through qualitative changes in αSyn structure or composition? If so, what defines seed-competent αSyn species in the human brain, and how do these properties influence their ability to template native αSyn, propagate pathology, and trigger cellular dysfunction? Moreover, are the seeds that spread through neural circuits the same species that impair mitochondria and lysosomes? And what molecular pathways mediate this transformation from a benign monomer into a self-perpetuating, organelle-disrupting aggregate?

In this study, we address these foundational questions using an integrated, multi-scale approach. By combining quantitative modeling of αSyn aggregation kinetics, cellular models derived from both sporadic and genetic PD cases, postmortem tissue spanning early- and late-stage brain regions, we dissect how αSyn structure and composition govern seeding activity, cellular toxicity, and Lewy pathology. This framework bridges molecular properties with functional outcomes and clinical relevance, offering mechanistic insight into the dynamic mechanisms that drive disease progression.

## Results

### Mitochondrial dysfunction in patient-derived neuron-like cells (iNLs)

To investigate the cellular mechanisms underlying PD, we used induced neuron-like cells (iNLs). These cells are previously generated by directly converting patient fibroblasts into induced neuronal progenitor cells (iNPCs), a process that preserves age-associated signatures and captures disease-relevant features^29,30^. We employed iNLs from four PD backgrounds, one with an *SNCA* triplication, two sporadic PD cases (sPD1 and sPD2), and one with a *PRKN* mutation, each paired with an age-matched neurologically healthy control (NHC). These lines have previously been used to model both genetic^31^ and sporadic^32^ forms of PD. This approach allowed us to study how αSyn-related alterations, including aberrant aggregation and post-translational modifications, drive mitochondrial dysfunction, a central feature of both familial and sporadic PD pathogenesis^32–34^. For all lines, the iNPCs expressed characteristic neural progenitor markers Pax6 and Nestin (**Figures S1A-1C**), while the resulting iNLs expressed neuronal marker Tuj1 and dopaminergic markers dopamine transporter (DAT), and tyrosine hydroxylase (TH) (**Figure 1C**). Consistent with prior reports, fibroblasts showed no detectable αSyn (**Figure S2**), while αSyn expression was induced in all iNLs, regardless of genetic background or disease status.

**Figure 1.**
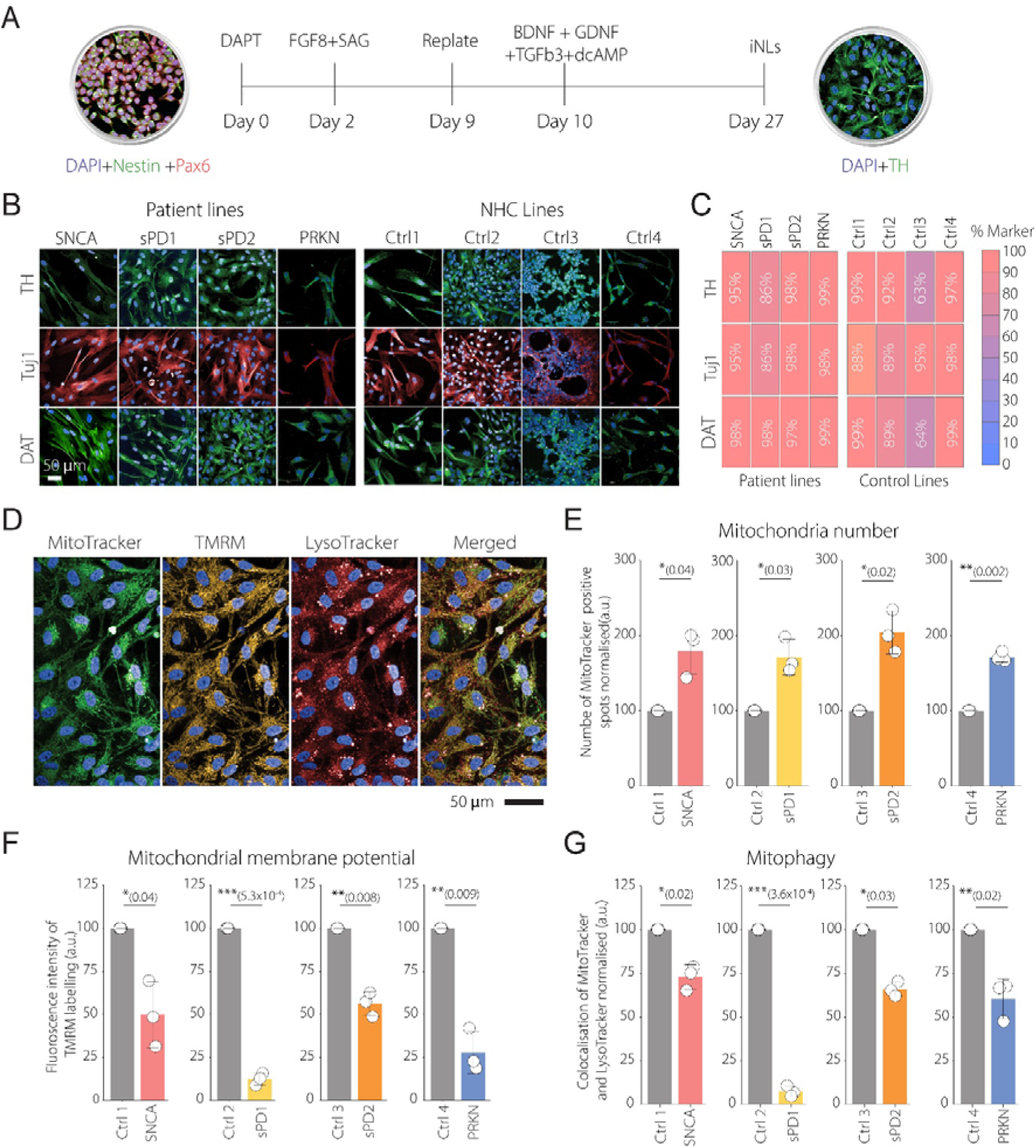
Assessment of mitochondrial quality control in PD and NHC iNLs. **(A)** Schematic of the preparation of iNLs from induced neural progenitor cells (iNPCs). **(B)** Immunofluorescence staining of iNLs for dopaminergic and neuronal markers: TH, Tuj1, and DAT in patient and age-matched control lines. **(C)** Quantification of marker-positive cells as a percentage of total nuclei (Hoechst) **(D)** Representative live-cell imaging of iNLs labelled with MitoTracker, TMRM, and LysoTracker. **(D)** Quantification of MitoTracker-positive puncta, normalized to total cell number per field of view. **(E)** Quantification of TMRM fluorescence intensity in MitoTracker-positive puncta, reflecting mitochondrial membrane potential and normalized to the respective control line. **(F)** Colocalization of MitoTracker and LysoTracker signals used as a proxy for mitophagy, normalized to respective control values Data are mean ± SD from biological replicates, with each replicate representing a separate neuronal differentiation. Statistical significance was determined by unpaired two-sample t-test. *P ≤ 0.05, **P ≤ 0.01, ***P ≤ 0.001; ****P ≤ 0.0001; ns not significant (P ≥ 0.05).

To assess mitochondrial function, we performed three-channel live-cell imaging with MitoTracker, TMRM, and LysoTracker across hundreds of iNLs from multiple differentiations (**Figure 1D**). Patient-derived iNLs exhibited increased mitochondrial number (**Figure 1E**), reduced mitochondrial membrane potential (**Figure 1F)** and decreased MitoTracker-LysoTracker colocalization, indicative of impaired mitophagy (**Figure 1G**). These findings suggest that mitochondrial dysfunction and disrupted quality control are shared phenotypes in our iNLs from both familial and sporadic PD patients, aligned with previous results^31,32^.

### Relative aggregation burden of αSyn drives LBs like pathology in PD iNLs

Given the consistent abnormalities across all patient-derived iNLs, we next asked whether specific αSyn species were associated with these mitochondrial defects (**Figure 2A**). To address this question, we quantified total wild-type (WT) αSyn (**Figure 2B**), its phosphorylated form (**Figure 2C**), and aggregated αSyn (**Figure 2D**) in cell lysates using a Meso Scale Discovery assay. Due to high variability in αSyn expression among individual control iNLs, we pooled data from four NHC lines to create a unified baseline group for comparison.

**Figure 2.**
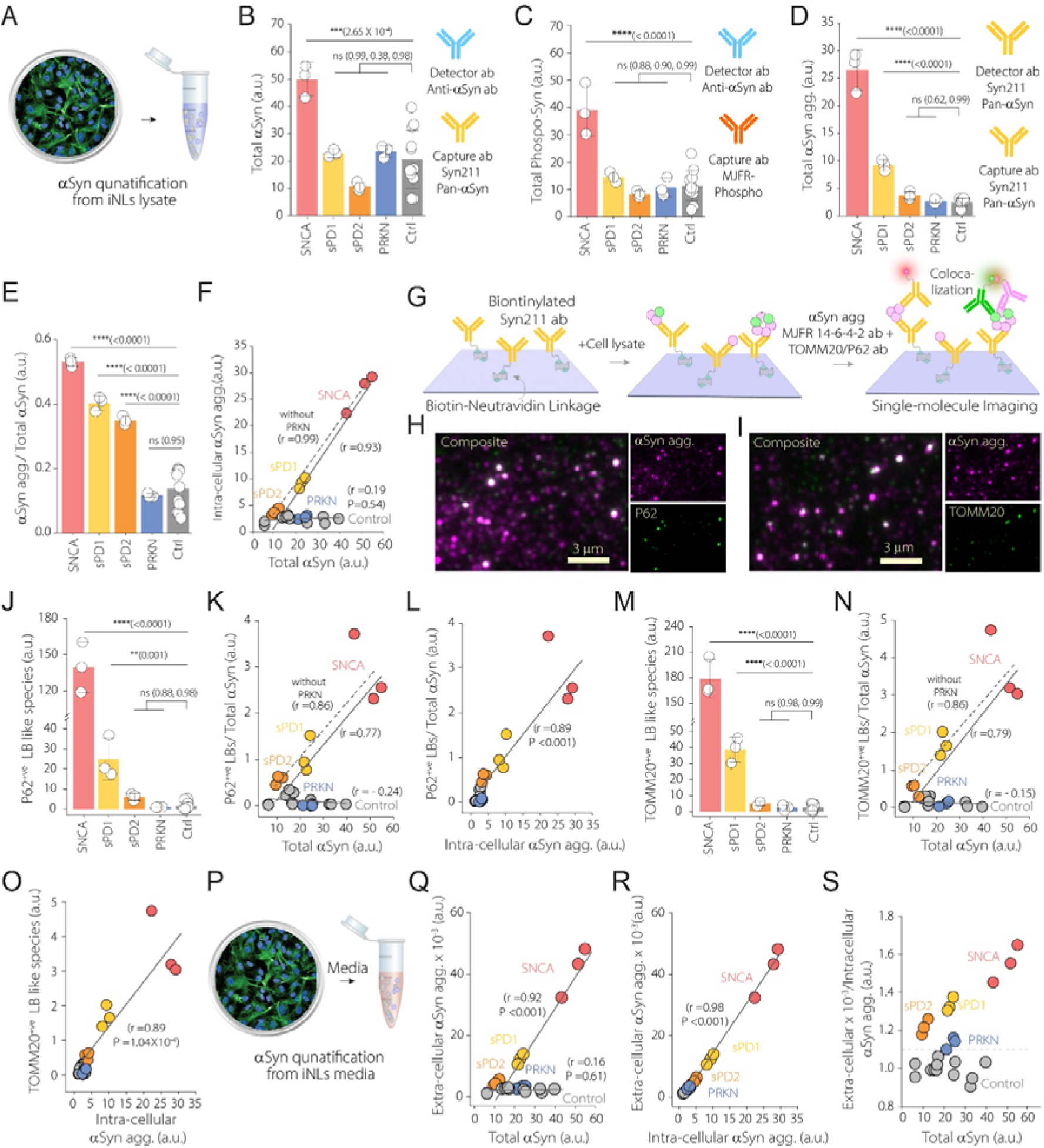
Aggregation and colocalization of α-syn with LB markers in iNLs. **(A)** Schematic of intracellular αSyn quantification from cell lysate **(B-D)** Quantification of (B) total αSyn, (C) phosphorylated αSyn (pS129), and (D) aggregated αSyn in lysate across cell lines using a MSD assay. (**E**) Ratio of aggregated αSyn to total αSyn. (**F**) Correlation between intracellular αSyn aggregates and total αSyn level across lines. (**G**) Schematic of the two-color SIMPull assay used to detect colocalization between αSyn aggregate preferring MJFR-14-6-4-2 antibody and with TOMM20 or p62 specific antibody (**H-I**) Representative SiMPull images showing colocalization of αSyn aggregates with p62 (H) or TOMM20 (I). (**J, M**) Quantification of αSyn aggregates colocalizing with p62 (J) or TOMM20 (M). (**K, N**) Correlation between total αSyn level and its colocalization with p62 (K) or TOMM20 (N). (**L, O**) Correlation between intracellular αSyn aggregates and its colocalization with p62 (L) or TOMM20 (O) across different cell lines. (**P**) Schematic of extracellular aggregate quantification from iNL conditioned media. (**Q**) Correlation between extracellular αSyn aggregates and total αSyn. (**R**) Correlation between extracellular and intracellular αSyn aggregates. (**S**) Extracellular-to-intracellular αSyn aggregate ratio plotted against total αSyn level. All measurements were performed on total protein normalized lysates (BCA assay). Data are mean ± SD from biological replicates, with each replicate representing a separate neuronal differentiation. Significance was determined by one-way ANOVA with post hoc Tukey-tests; correlations used Pearson’s r. *P ≤ 0.05, **P ≤ 0.01, ***P ≤ 0.001; ****P ≤ 0.0001; ns not significant (P ≥ 0.05).

As expected, the *SNCA* triplication line exhibited the highest levels of total WT αSyn and pS129 αSyn and aggregated αSyn. Total and pS129 αSyn levels in sPD1, sPD2, and *PRKN* lines were similar to controls. However, only sPD1, along with the *SNCA* line, exhibited elevated intracellular aggregates, while sPD2 and *PRKN* lines did not. To assess whether the aggregation burden was disproportionately high relative to total αSyn expression, we normalized αSyn aggregate levels to total αSyn for each line (**Figure 2E**). This revealed a significantly elevated aggregate-to-total αSyn ratio in the *SNCA* and both sporadic PD lines, but not in the *PRKN* line.

To further explore the relationship, we examined the correlation between αSyn aggregates and total αSyn levels across all iNLs (**Figure 2F**). In NHC lines, no significant correlation was observed (r = 0.19, P = 0.54), suggesting αSyn burden alone does not drive aggregation. In contrast, patient lines showed a strong positive correlation between total αSyn and aggregate levels (r = 0.93, P < 0.0001 including *PRKN*; r = 0.99, P < 0.0001 excluding *PRKN*), indicating that elevated αSyn level in the disease context is closely associated with increased aggregation.

To investigate whether these aggregates are associated with common dysfunctions implicated in PD, we employed a <u>si</u>ngle-<u>m</u>olecule <u>pull</u>-down (SiMPull) assay^35^ to investigate the association of αSyn aggregates with hallmark LB components and organelles implicated in PD pathology, directly from cell lysates without prior purification (**Figure 2G**). Aggregated αSyn was captured using the conformation-selective MJFR-14-6-4-2 antibody and assessed for colocalization with canonical LB markers, including phosphorylated αSyn (pS129), the autophagy adaptor p62, ubiquitin, and lipid components, as well as mitochondrial (TOMM20, VDAC1) and lysosomal (LAMP2) proteins (**Figure 2H-I, S3A-3F).** Although TOMM20, VDAC1, and LAMP2 are not LB markers, their association with αSyn aggregates indicates mitochondrial and lysosomal involvement, two organelles whose dysfunction is known to contribute to LB formation and PD pathogenesis. For the remainder of the study, we focus on two representative markers: p62, a canonical component of Lewy pathology, and TOMM20, which indicates mitochondrial interaction with αSyn aggregates.

The *SNCA* triplication line and sPD1 showed significantly increased colocalization of αSyn aggregates with both p62 (**Figure 2J**) and TOMM20 (**Figure 2M**) compared to NHCs, whereas sPD2 and the *PRKN* line did not. Correlation analysis showed that p62 (**Figure 2K**) and TOMM20 (**Figure 2N**) levels were correlated with total αSyn level only in patient lines (p62: r = 0.86 with *PRKN*, r = 0.77 without; TOMM20: r = 0.86 with *PRKN*, r = 0.79 without), whereas only a weak link was observed in control lines (p62: r = 0.24; TOMM20: r = 0.15) (Figure 2K, 2N). Importantly, intracellular αSyn aggregate levels were positively correlated with LB-like species marked by p62 (r = 0.93, P < 0.0001) (**Figure 2L**) and TOMM20 (r = 0.95, P < 0.0001) (**Figure 2O**) colocalization, indicating that spontaneous LB-like formation in iNLs is tightly linked to αSyn aggregation burden, independent of genetic background.

### Intracellular αSyn aggregation drives extracellular αSyn release

We next quantified extracellular αSyn aggregates in the conditioned media from iNLs (**Figure 2P**). In control lines, extracellular aggregates did not correlate significantly with total αSyn level (r = 0.16, P = 0.61). In contrast, a strong positive correlation was observed in patient-derived iNLs (r = 0.93, P < 0.0001) **(Figure 2Q**). We also found a strong link between intracellular and extracellular aggregate levels across all lines (r = 0.98, P < 0.0001) (**Figure 2R**), suggesting that presence of increased intracellular aggregation drives extracellular αSyn release. We then calculated the ratio of extracellular to intracellular aggregates for each line (**Figure 2S**). In control iNLs, this ratio stayed relatively constant, regardless of αSyn levels. However, in PD iNLs, the ratio was consistently higher and increased further with more αSyn expression. These results suggest that PD iNLs release αSyn aggregates at much higher rates than controls, especially when total αSyn levels are elevated.

### αSyn fibrils induces mitochondrial dysfunction and LB like species formation in iNLs

Given the elevated presence of extracellular αSyn aggregates in PD iNLs, we hypothesized that these species may act as pathological seeds - a key mechanism implicated in disease progression. To systematically evaluate how different aggregation state (oligomer vs. fibril) and phosphorylation influence the seeding activity, we generated αSyn aggregates. Monomeric WT αSyn was first phosphorylated using PLK3^36^, resulting in phosphorylation at serine 129 (pS129) as well as additional serine (S42, S87) and tyrosine (T33, T64, T92) residues, confirmed by mass spectrometry (**Figure S4**). Phosphorylation was also validated by western blot using pan-αSyn and pS129-specific antibodies (**Figure 3A**). Both monomeric WT and phosphorylated αSyn (70 μM) were then aggregated as previously mentioned^33,37^, and their aggregation was monitored using thioflavin T (ThT) fluorescence, which revealed comparable rates for both forms (**Figure 3B**). Single-aggregate imaging^38^ showed that the samples close to the lag phase contained in small, circular oligomer like intermediates, whereas in the plateau phase, the samples contained mostly elongated, fibrillar species (**Figure 3C**).

**Figure 3.**
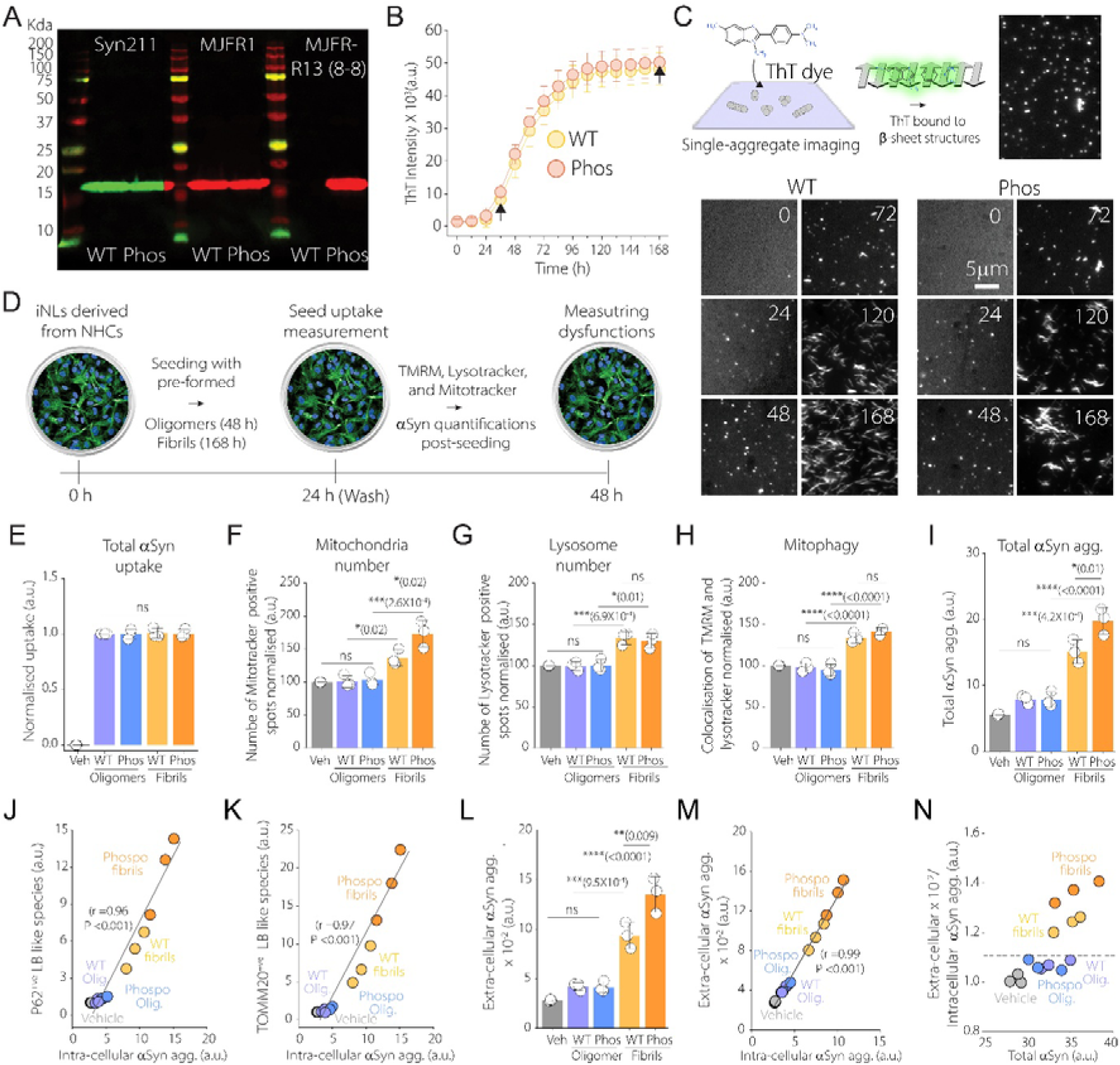
Assessing the impact of WT and phosphorylated αSyn seeds on αSyn aggregation, LB-like species formation, and mitochondrial function in iNLs. **(A)** Western blot confirming phosphorylation of recombinant αSyn. Total αSyn was detected using Syn211 and MJFR1 antibodies, and phosphorylated αSyn was detected using the phospho-specific MJF-R13 (8-8) antibody. **(B)** Aggregation kinetics of WT and phosphorylated αSyn measured using ThT, with selected time points for downstream analysis indicated by black arrows **(C)** Schematic of single-aggregate imaging using ThT and representative images of ThT-stained aggregates at different time points during the aggregation for WT and phosphorylated αSyn. **(D)** Schematic of the experimental timeline iNLs based seeding assay. **(E)** Quantification of Alexa Fluor 488 dye labelled αSyn as measure of uptake across treatment groups **(F-G)** Mitochondrial and Lysosomal number measured using TMRM (F) and Lysotracker (G) respectively, both normalised to cell number. **(H)** Mitophagy levels quantified by colocalization of TMRM and LysoTracker, normalised to cell area **(I)** Total intracellular αSyn aggregate measured post-seeding. **(J-K)** Quantification of colocalization between αSyn aggregates with p62 (J) and TOMM20 (K). **(L)** Extracellular αSyn aggregates measured in conditioned media following different seed treatment. **(M)** Correlation between intracellular and extracellular αSyn aggregates across seed treatment. **(N)** Ratio of extracellular to intracellular αSyn aggregates with total αSyn for different conditions. Data are mean ± SD from biological replicates, with each replicate representing a separate neuronal differentiation. Statistical significance was determined by one-way ANOVA with post hoc Tukey-tests; correlations used Pearson’s r. p-values: *P ≤ 0.05, **P ≤ 0.01, ***P ≤ 0.001; ****P ≤ 0.0001; ns not significant (P ≥ 0.05).

These distinct αSyn aggregate preparations, oligomer-enriched samples from the end of the lag phase and fibrillar-enriched sonicated samples from the plateau phase, were generated from both wild-type and phosphorylated αSyn. They were added separately to NHC-derived iNLs at a final concentration corresponding to 1 µM monomer equivalent (**Figure 3D).** Although aggregation reactions at any time point yield heterogeneous mixtures of unreacted monomers, soluble oligomers, and fibrils, these two time points were selected to represent stages enriched in oligomeric intermediate and mature fibril species, respectively, based on their aggregation kinetics and morphology. Cells were treated with these aggregate preparations for 24 hours, followed by a media change and an additional 24-hour incubation to assess downstream effects.

To confirm that any cellular responses were not due to differences in uptake, we first measured internalization of Alexa Fluor 488 (AF488) labelled αSyn and its phosphorylated forms. Phosphorylation of labelled αSyn was verified by western blot (**Figure S5**). Fluorescent labelling allowed us to distinguish exogenous aggregates which are used as seed from endogenous αSyn. Quantification of dye intensity in iNLs showed no significant differences in uptake among different seeds regardless of aggregation state or phosphorylation status (**Figure 3E**) and confirms that any observed differences in downstream effects are not attributable to variations in seed internalization.

We then assessed how these different αSyn seeds influence mitochondrial function. Seeds isolated from end of the lag phase of the aggregation reactions, containing an oligomer population, both wild-type (WT) and phosphorylated, did not alter mitochondrial number (**Figure 3F**), lysosomal number (**Figure 3G**), or mitophagy (**Figure 3H**). In contrast, both WT and phosphorylated αSyn fibrils caused a significant increase in mitochondrial and lysosomal numbers, along with enhanced mitophagy. Given that mitophagy is typically reduced in PD iNLs, this increase upon seeding reflects a compensatory response to the sudden influx of fibrillar seeds, triggering mitochondrial stress and prompting the cell to enhance mitophagy as a means to restore homeostasis.

We then determined whether the observed dysfunctions were linked to the extent of intracellular αSyn aggregation following seeding. Aggregation mixtures containing oligomers did not increase aggregate burden relative to unseeded controls, whereas both wild-type and phosphorylated fibrils triggered substantial aggregation, with phosphorylated fibrils inducing the highest levels (**Figure 3I**). These aggregates colocalized strongly with p62 (**Figure 3J**) and TOMM20 (**Figure 3K**), and their abundance closely correlated with the number of nanoscopic αSyn species positive for p62 (r = 0.96, P < 0.0001) and TOMM20 (r = 0.97, P < 0.0001).

We also quantified extracellular αSyn aggregates, as prior studies have suggested that LB formation can be promoted by extracellular αSyn^22,23^. Neither WT nor phosphorylated seeds isolated from earlier stages of aggregation led to a significant increase in extracellular αSyn levels. In contrast, both WT and phosphorylated fibrillar seeds significantly elevated extracellular αSyn, with phosphorylated fibrils inducing the highest levels **(Figure 3L**). This mirrored the intracellular aggregation pattern, and we observed a strong positive correlation (r = 0.99, P < 0.0001) between intracellular and extracellular αSyn aggregates (**Figure 3M**), suggesting that increased intracellular seeding drive αSyn release into the extracellular space. Finally, we assessed the ratio of extracellular to intracellular aggregates. While this ratio remained unchanged following oligomer containing sample treatment, both WT and phosphorylated fibrils caused a disproportionate increase in extracellular αSyn (**Figure 3N**), further supporting the idea that only fibrillar seeding drives both intracellular LB-like species formation and extracellular aggregate release.

### Fibrillar αSyn seeds promote damaging aggregates formation via secondary nucleation

Given our observation that fibrillar αSyn aggregates, both WT and phosphorylated, induce mitochondrial dysfunction and LB like pathology in iNLs derived from NHCs, we next sought to uncover the mechanistic basis for this difference. To isolate the intrinsic effects of these seeds from confounding cellular processes such as uptake, trafficking, or degradation, we employed a recombinant αSyn-based *in vitro* system. This system allowed us to determine whether the differential toxicity stems from differences in the ability of fibrils versus oligomers to accelerate αSyn aggregation and generate new, damaging species.

For this combined assay, pre-formed WT or phosphorylated αSyn aggregates, either oligomer- or fibril-containing preparations, were added at varying concentrations (0.1%, 0.3%, and 1%) to 70 µM monomeric αSyn and aggregation was monitored using ThT (**Figure 4A**). In parallel, aliquots were taken at defined time points and tested for toxicity using a single-vesicle membrane permeabilization assay. We used membrane permeabilization as a measure of their damaging ability, given that αSyn aggregates toxicity has been widely associated with membrane perturbation^39–41^. Here, nanoscale lipid vesicles preloaded with a Ca²⁺-sensitive dye were exposed to reaction aliquots. Aggregates in the samples permeabilized the vesicles, allowing Ca²⁺ influx and resulting in increased fluorescence^42^. Each vesicle acts as an independent reaction chamber, providing single-molecule sensitivity and allowing the detection of damaging αSyn species even in complex biological systems such as cerebrospinal fluid^43^, stem cell-derived neurons^33^, and post-mortem brain tissue^44^.

**Figure 4.**
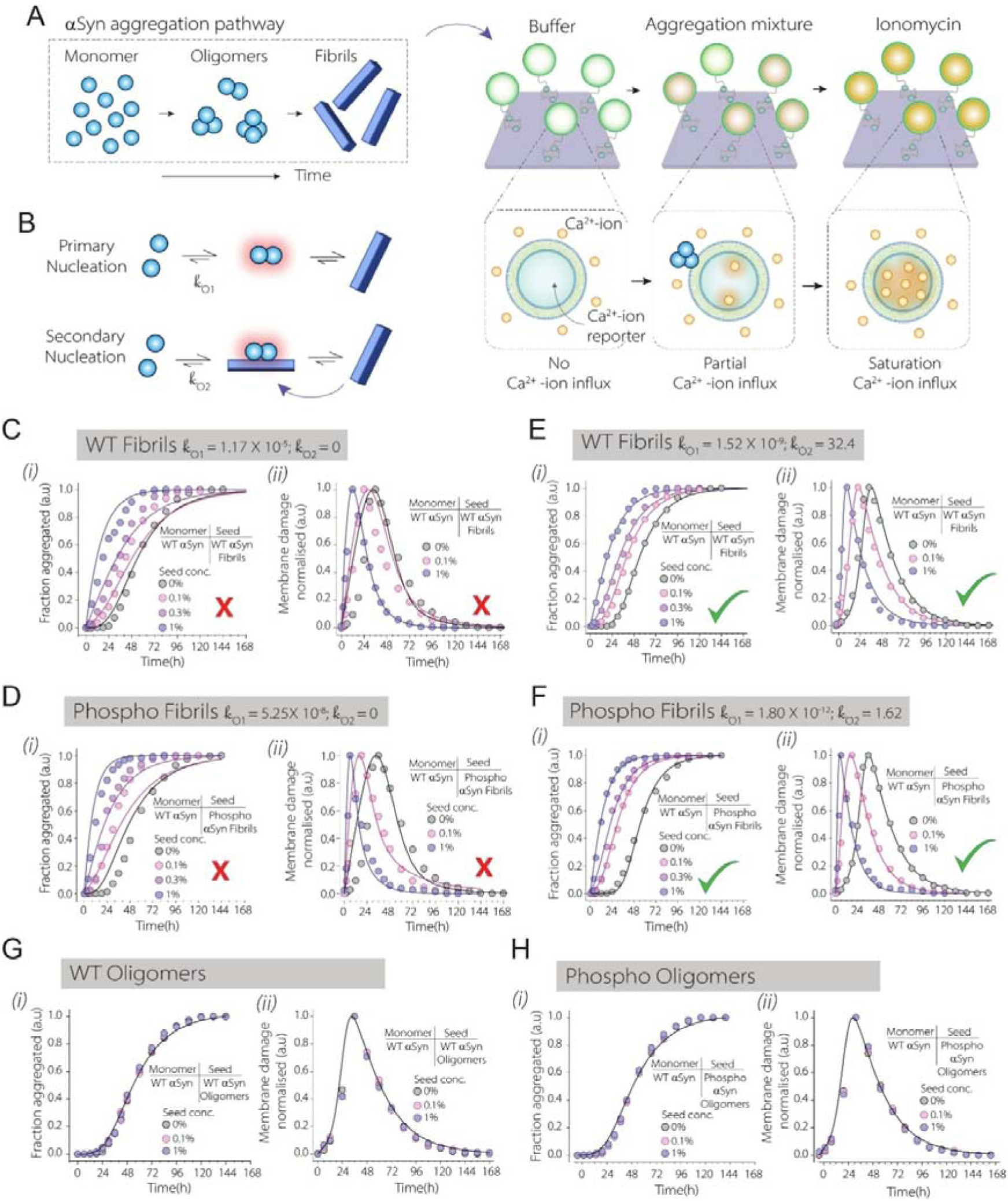
Relationship betweenαSyn pre-formed seed type, aggregation kinetics, and damaging ability of newly formed aggregates. **(A)** Schematic of the experimental workflow. Aggregation pathways and membrane-disrupting potential of newly formed aggregates were assessed by a membrane permeabilization assay. **(B)** Schematic illustrating primary and secondary nucleation pathways **(C-F)** Aggregation kinetics of monomeric αSyn *(i)* and corresponding membrane permeabilization profiles *(ii),* in the presence of WT (C, E) or phosphorylated (D, F) fibrils. (C, D) Data fitted using a model with primary nucleation only (k_₂_ = 0).(E, F) Data fitted using a combined model including both primary and secondary nucleation pathways. **(G-H)** Aggregation kinetics *(i)* and membrane permeabilization profiles *(ii)* in the presence of WT (G) or phosphorylated (H) oligomers. No differences were observed compared to unseeded monomeric aggregation. Experimental data points are shown as circles, and lines represent the kinetic fitting.

Using this combination assay, we observed that membrane-disrupting activity was absent at the onset of aggregation, peaked near the end of the lag phase, and gradually declined before diminishing completely during the plateau phase. This temporal pattern suggests that membrane damage is primarily mediated by transient species, such as oligomeric intermediates, generated during early stages of the fibrillation pathway. The pronounced peak in damaging activity at the end of the lag phase for both WT and phosphorylated αSyn suggests that this stage represents the point of highest toxic oligomer abundance across the entire aggregation process. Concurrently, ThT fluorescence was minimal at this point, indicating a low level of β-sheet–rich fibrils and supporting our selection of this time point as the source of oligomer-enriched preparations. In contrast, samples collected during the plateau phase exhibited maximal ThT fluorescence, consistent with abundant fibril formation, but showed negligible membrane-disrupting activity, further validating their use as fibrillar seed preparations. Since both sample sets were derived from the same initial monomer concentration and differ only in the timing of collection, this experimental design enables a reasonable and controlled comparison of the seeding activity of oligomeric intermediates versus fibrils, despite the inherent heterogeneity of each sample.

To determine the pathway responsible for generating these damaging intermediates during the fibrillation pathway, we fitted both ThT fluorescence and membrane permeabilization data to a kinetic model incorporating two nucleation mechanisms^45^: *(i)* primary nucleation, where monomers spontaneously form aggregates, and *(ii)* secondary nucleation, where monomers aggregate on the surface of existing pre-formed aggregates (**Figure 4B**). When the model was constrained to primary nucleation only (k₂ = 0), it failed to recapitulate the experimental data observed for both WT (**Figure 4C**) and phosphorylated αSyn fibrils (**Figure 4D**). In contrast, including secondary nucleation in the model provided an excellent fit to the data (**Figure 4E, F**), indicating that fibrillar surfaces serve as highly efficient catalytic platforms for generating new, potentially damaging aggregates. This mechanism which was recently recognized as the dominant route of αSyn oligomer generation, is considerably more efficient than either primary nucleation or fibril fragmentation^46^.

In contrast, seeding with oligomer-containing preparations failed to accelerate aggregation or enhance membrane permeabilization under equivalent conditions for either WT (**Figure 4G**) or phosphorylated αSyn (**Figure 4H**). The absence of enhanced aggregation or membrane disruption from peak oligomer samples suggests that fibrillar αSyn seeds are more potent in driving toxic intermediate formation through secondary nucleation.

### Phosphorylated αSyn aggregates resist degradation and disrupt mitochondrial membranes

While both WT and phosphorylated αSyn fibrils effectively seed aggregation, phosphorylated fibrils consistently caused greater dysfunction in control iNLs. To investigate whether differences in their susceptibility to cellular degradation could contribute to this disparity, we studied their susceptibility to two key clearance pathways: the autophagy-lysosome pathway (ALP) and the ubiquitin-proteasome pathway (UPP). To assess the role of each pathways in αSyn aggregate clearance following seeding, iNLs derived from a NHC were pre-treated with chloroquine (CQ) to inhibit ALP or MG132 to inhibit the UPP, followed by seeding with buffer (unseeded), WT αSyn fibrils, or phosphorylated αSyn fibrils. Intracellular αSyn aggregate was quantified 8h later (**Figure 5A**). CQ treatment significantly increased αSyn aggregates in unseeded and WT-seeded cells, while only a modest, non-significant increase was observed with phosphorylated fibrils (**Figure 5B**). The fold increase was highest in the unseeded condition, followed by WT, and lowest for phosphorylated αSyn fibrils (**Figure 5C**). In contrast, inhibition of UPP function led to an increase in αSyn aggregates across all conditions, including unseeded, WT-seeded, and phosphorylated fibril-seeded cells (**Figure 5D**). However, the relative fold changes in aggregation were similar between WT and phosphorylated fibril conditions (**Figure 5E**), indicating that the UPP does not differentially process these fibril species.

**Figure 5.**
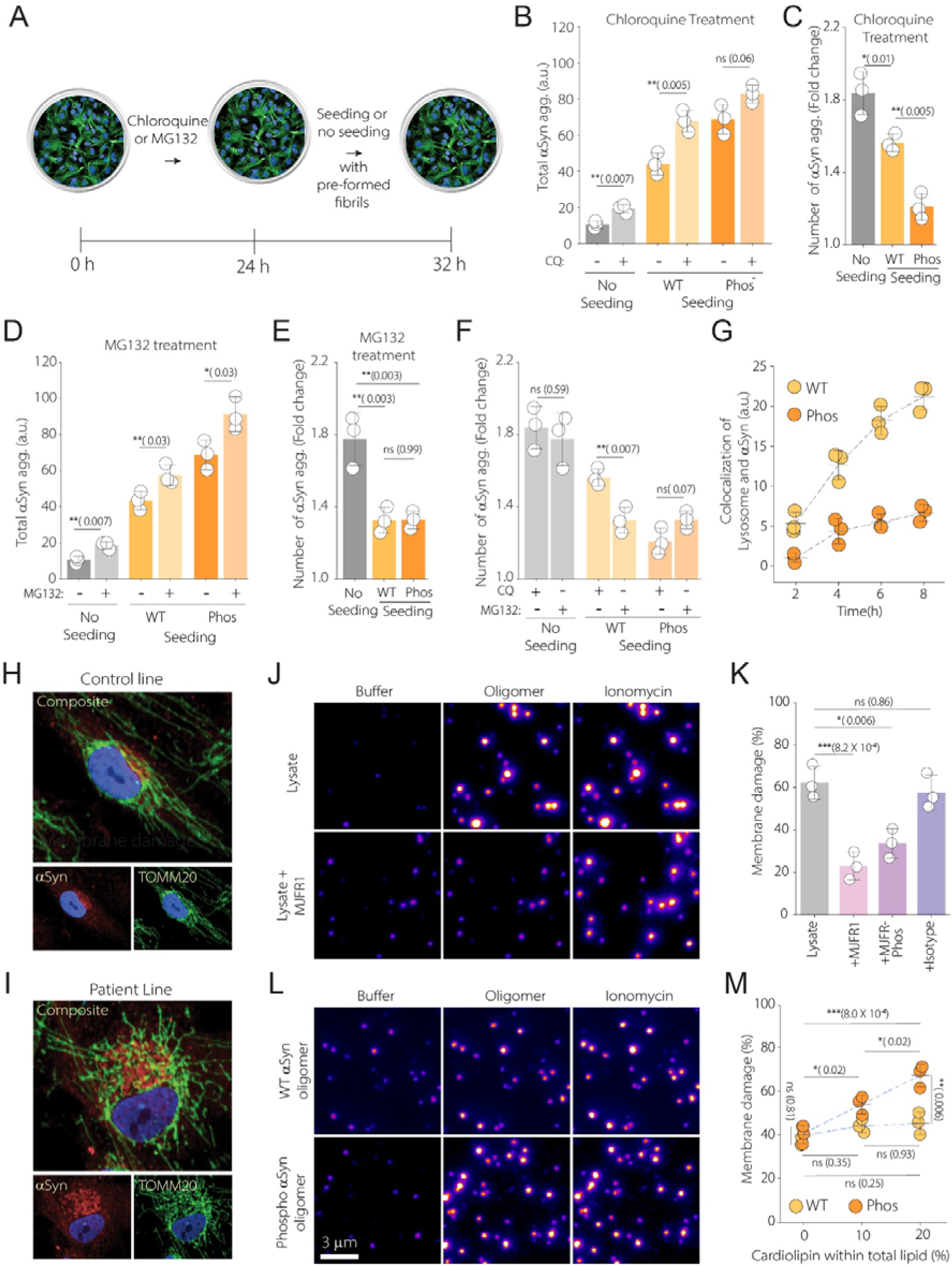
Phosphorylated αSyn evade autophagy-lysosomal clearance and disrupt cardiolipin-rich membranes. **(A)** Schematic of the experimental timeline showing pharmacological inhibition and seeding in iNLs. **(B, C)** Quantification of intracellular αSyn aggregate s(B) and fold change after ALP inhibition by chloroquine with or without seeding (C). **(D, E)** Quantification of intracellular αSyn aggregates **(D)** and fold change after UPP inhibition by MG132 with or without seeding (E). **(F)** Comparison of fold change in number of αSyn aggregates between CQ and MG132 treatment with or without WT/Phos seeding. **(G)** Time-lapse live-cell imaging showing colocalization of Alexa Fluor 488 dye labelled WT and phospho αSyn fibrils and LysoTracker. **(H-I)** Immunofluorescence imaging of αSyn and TOMM20 in control (H) and patient-derived iNLs (I). **(J-K)** Representative images of membrane permeabilization assay when lipid vesicles incubated with iNLs-derived lysates from *SNCA* triplication line, either directly or following IP with MJFR1, MJF-R13 (8-8), or isotype control (J) and their quantification(K) **(L-M)** Representative images (20% cardiolipin) of membrane permeabilization assay using 100 nM recombinant WT and phosphorylated αSyn oligomers (L) and their quantification (M). Data are mean ± SD from biological replicates, with each replicate representing a separate neuronal differentiation (B-G). Statistical significance was assessed using unpaired two-sample t-tests (B, D, F) or one-way ANOVAs with post hoc Tukey’s tests (C, E, M, K); P-values are indicated as follows: *P < 0.05, **P < 0.01, *P < 0.001; ns, not significant (P ≥ 0.05).

To compare the roles of ALP and UPP in αSyn clearance, we measured aggregate levels after inhibiting each pathway. In unseeded and phosphorylated fibril-seeded cells, ALP and UPP inhibition caused similar increases. In contrast, ALP inhibition led to a much larger increase in aggregation in WT-seeded cells, indicating preferential clearance of post WT fibril seeding via the ALP (**Figure 5F)**. To investigate this, we used live-cell imaging to track lysosomal targeting of AF488 dye labelled WT and phosphorylated αSyn fibrils in iNLs stained with LysoTracker (**Figure 5G**). WT fibrils showed greater and more sustained colocalization with lysosomes than phosphorylated fibrils, indicating more efficient autophagic targeting. These findings suggest that phosphorylated fibrils, by evading lysosomal degradation, persist longer inside cells, leading to greater intracellular accumulation and disruption of cellular homeostasis.

We next investigated how WT and phosphorylated αSyn aggregates interact with mitochondria-like membranes, given the known association of αSyn with mitochondria^32,33^ (**Figure 5H,I**) and its involvement in LB formation^10^. To determine whether endogenous αSyn species in iNLs contribute to mitochondrial membrane disruption, we tested lysates from *SNCA*-overexpressing iNLs for their ability to permeabilize cardiolipin (20%)-containing vesicles. Cardiolipin is a mitochondria-specific phospholipid essential for maintaining mitochondrial bioenergetics^47^, and it has been shown to interact and modulate the pore-forming capacity of αSyn oligomers^48,49^, thereby contributing to mitochondrial dysfunction These lysates exhibited clear membrane-permeabilizing activity (**Figure 5K**). When the lysates were incubated with antibodies specific to total αSyn (MJFR1) or phosphorylated pS129 αSyn (MJF-R13 (8-8)) and then added to the vesicles, the membrane-disrupting activity was significantly reduced. This indicates that both WT and phosphorylated αSyn oligomers present in the lysates contribute to mitochondrial membrane damage.

To further determine the influence of cardiolipin on αSyn-membrane interactions, we prepared nanosized (Cut-off 100 nm) lipid vesicles containing 0%, 10%, or 20% cardiolipin to mimic mitochondrial membranes. We then assessed the membrane-disrupting ability of oligomeric aggregates generated in vitro from recombinant WT or phosphorylated αSyn (**Figure 5L**). In cardiolipin-free vesicles, there was no significant difference in membrane permeabilization between WT and phosphorylated αSyn oligomers. However, in cardiolipin-containing vesicles, phosphorylated αSyn oligomers induced significantly more membrane disruption in a cardiolipin concentration-dependent manner. In contrast, WT αSyn oligomers showed little to no change in membrane-permeabilizing activity across different cardiolipin concentrations (**Figure 5M**). These results also indicate that phosphorylated αSyn oligomers interact more strongly with cardiolipin-rich mitochondria mimicking membranes, enhancing their ability to disrupt membrane integrity.

### αSyn seeding activity in postmortem PD brain tissue correlates with the spatiotemporal progression and regional burden of LBs

To investigate how the αSyn dysfunction and seeding activity observed in our cellular model systems are reflected in human PD, we analysed postmortem brain tissue from five individuals with sporadic PD and five matched NHCs. For each individual, three brain regions were studied: the substantia nigra (SN), cingulate gyrus (CG), and neocortex (NC). These regions were selected based on the established spatiotemporal pattern of PD pathology, in which the SN is affected in earlier disease stages, followed by the CG and later the NC^19,20^. Formalin-fixed sections from each region were stained with an antibody against pSer129 αSyn to visualize LB by immunohistochemistry (**Figure 6A**). LB counts were significantly higher in PD cases compared to NHCs (**Figure 6B**). LB pathology in PD brains followed a region-specific distribution, with the SN showing the highest burden, followed by the CG and NC (**Figure 6C**), consistent with spatiotemporal pattern of PD progression. To assess whether total αSyn levels explain regional pathology, we quantified αSyn in frozen sections from the same brain regions. We found no correlation between total αSyn and LB burden (r = –0.22, P = 0.41; **Figure 6D**), consistent with our data form iNL models. This finding not only highlights that LB formation is not solely determined by αSyn abundance, but also validates our cell system as a relevant model for sporadic PD.

**Figure 6.**
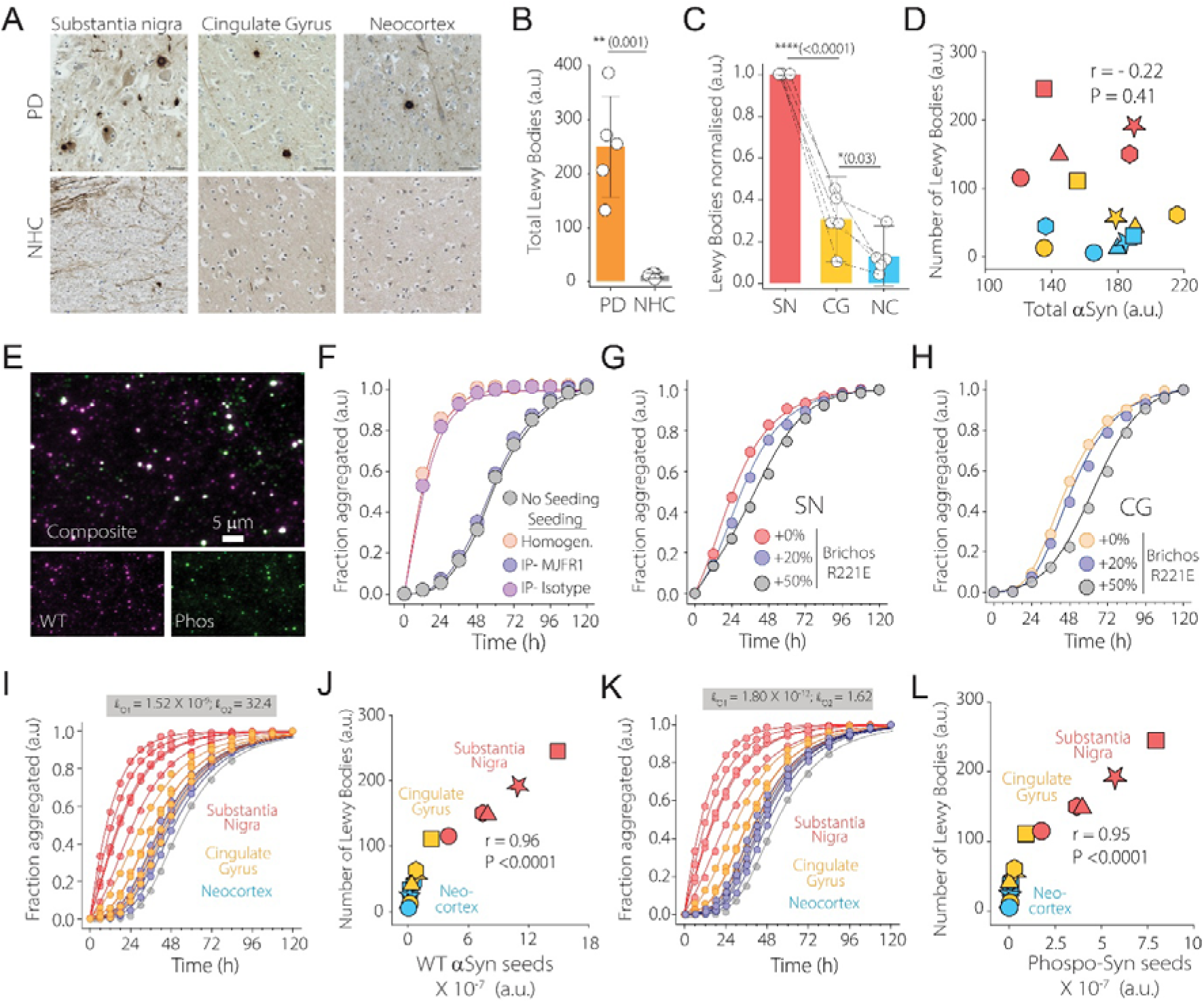
Correlation ofαSyn seeding activity and LB pathology in postmortem PD brain tissue. **(A)** Representative pSer129-αSyn immunohistochemistry images from substantia nigra, cingulate gyrus, and neocortex of PD patients and NHCs. **(B–C)** Quantification of total LBs (B) and their distribution across brain regions (C). **(D)** Correlation of total αSyn levels and LB counts across regions. Shape of the symbols indicates patient, and colour indicates brain region. **(E)** SiMPull imaging of WT and phosphorylated αSyn species in SN homogenate. **(F)** Aggregation of recombinant αSyn seeded with PD brain homogenate, with or without immunodepleting using MJFR1 or isotype control antibodies. **(G–H)** Aggregation in the presence of Bri2 BRICHOS R221E (20% or 50%) to assess inhibition of αSyn secondary nucleation when seeded with homogenates from the substantia nigra (G) or cingulate gyrus (H). Circles represent experimental data; lines show model fits. **(I-L)** Representative aggregation of recombinant αSyn seeded with homogenates from SN, CG, and NC, fitted using WT (I) and phosphorylated αSyn (K) kinetic parameters. Correlation of estimated WT (J) and phosphorylated (L) αSyn seed concentrations with LB counts from matched brain regions. Statistical significance was assessed using unpaired two-sample t-tests (B) or one-way ANOVAs with post hoc Tukey-tests (C); p-values are indicated as follows: *P < 0.05, **P < 0.01, *P < 0.001; ns, not significant (P ≥ 0.05).

We next assessed whether αSyn seeds correlate more closely with Lewy pathology. To first confirm the presence of αSyn aggregates, we performed SiMPull imaging on SN homogenates from a PD brain. This revealed both wild-type and phosphorylated αSyn species, with partial colocalization (**Figure 6E**). To determine whether these species possess seeding ability, we incubated the homogenates with recombinant monomeric αSyn. Compared to unseeded controls, the homogenates markedly accelerated aggregation, confirming the presence of seeding-competent species in PD brain tissue (**Figure 6F**). Immunodepletion using the αSyn-specific MJFR1 antibody significantly reduced seeding activity, demonstrating that αSyn-positive aggregates are the primary drivers of seeding in these homogenates (**Figure 6F**). To test whether this activity involves secondary nucleation, we used recombinant Bri2 BRICHOS R221E, a chaperone known to selectively inhibit the secondary nucleation pathway^50,51^. Recombinant αSyn was seeded with either SN (**Figure 6G**) or CG (**Figure 6H**) homogenates in the presence of increasing concentrations of Bri2 BRICHOS R221E monomer (20% or 50%). We observed a dose-dependent reduction in secondary nucleation without changes in primary nucleation or elongation rates. These findings suggest that αSyn seeds from PD brain tissue homogenates accelerate monomer aggregation via secondary nucleation.

Finally, to estimate the abundance of seeding-competent species across different brain regions, we performed seeded aggregation assays using homogenates from all PD patients and brain regions. These kinetic data were then fitted using the quantitative framework we developed earlier. Since these homogenates contain both WT and phosphorylated αSyn species (**Figure 6E**), and their combined effects are difficult to deconvolve, we modelled each dataset separately under the assumption that seeds consisted entirely of either WT (**Figure 6I**) or phosphorylated αSyn (**Figure 6K**) seeds. When estimated seed concentrations were compared with LB counts from the corresponding regions, both WT (r = 0.95, P < 0.0001; **Figure 6J**) and phosphorylated αSyn (r = 0.96, P < 0.0001; **Figure 6L**) seed levels showed strong correlations with regional pathology.

## Discussion

The spread of Lewy pathology along anatomical networks is a defining hallmark of PD^19,20^, and cell-to-cell transmission of αSyn aggregates is considered a major driver of this process^21,22^. The diagnostic success of αSyn seed amplification assays^26–28^ lends further support to this hypothesis. However, how αSyn seeding activity translates into burden and distribution of Lewy pathology, and ultimately contributes to neuronal loss and PD progression, remains largely unresolved. Here, we bridge this critical gap by systematically mapping the molecular choreography of αSyn-seeded pathology, beginning with seed uptake and degradation (or evasion), through the engagement of endogenous αSyn, the formation of new aggregates, and the onset of organelle dysfunction, culminating in LB like species formation. Our results show that this pathological sequence is influenced by the structural and compositional features of the initiating αSyn seeds, which regulate both the efficiency of new aggregate generation and the severity of downstream cellular damage. Our framework predicts the spatiotemporal progression and regional burden of Lewy pathology in the human brain, establishing a direct mechanistic link between αSyn seeding, LB formation and the pathological staging of PD at the molecular scale.

We used directly reprogrammed iNLs as model system, as this model retains age-associated features and recapitulate key PD hallmarks^31,32^, including mitochondrial vulnerability and the spontaneous formation of nanoscopic LB-like species. Although morphologically distinct from the large, mature LBs observed in postmortem PD brain, these nanoscopic LB-like species consistently stain for canonical LB markers such as p62 and ubiquitin^10^. They also co-localize with organelle-associated proteins frequently implicated in PD pathogenesis, including the mitochondrial markers TOMM20 and VDAC1, and the lysosomal marker LAMP2. Notably, exposure of control iNLs to exogenous αSyn seeds was sufficient to induce LB-like inclusion formation and mitochondrial impairment, mirroring the pathological features seen in PD patient-derived lines. This consistent relationship between αSyn seeded aggregation, mitochondrial disruption, and LB-like pathology suggests that iNLs capture early, disease-relevant events in αSyn-mediated neurodegeneration, offering a tractable, physiologically grounded model system.

To benchmark this model across the pathological spectrum of PD, we incorporated two genetically distinct lines representing opposite ends of the αSyn pathology: an *SNCA* triplication line with high αSyn burden^4^ and a *PRKN* mutant line that lacks Lewy pathology despite neurodegeneration^7^. Given the convergence of multiple genetic forms of PD on shared mechanisms such as mitochondrial dysfunction^9^, these two lines served as contrasting reference points for interpreting αSyn-driven changes in idiopathic PD. In iNLs derived from sporadic PD patients, we observed an increased ratio of aggregated to total αSyn relative to controls, positioned between the *SNCA* and *PRKN* lines. This elevated aggregation burden correlated strongly with both the formation of LB-like inclusions and the accumulation of extracellular αSyn aggregates, indication of active intracellular seeding. To identify which αSyn assemblies most potent seed, we exposed control iNLs to oligomer- and fibril-enriched αSyn preparations generated from equivalent monomer concentrations. Strikingly, αSyn fibrils induced significantly more severe mitochondrial dysfunction and LB-like species formation.

This was unexpected, as αSyn oligomers have been reported as highly cytotoxic in the αSyn aggregation pathway^39,40^. To better understand this apparent contradiction, we examined how each type of species influenced intracellular αSyn aggregation. We observed that iNLs seeded with fibrils accumulated significantly more intracellular aggregates than those treated with samples collected at the aggregation time point corresponding to peak oligomer levels or left untreated. These findings suggest that the observed dysfunction might not result from the seeds themselves, but from their capacity to drive the formation of new, toxic species within cells. To test this possibility, we developed an integrated framework combining recombinant αSyn aggregation kinetics, membrane permeabilization assays to assess the damaging capacity of newly formed species, and theoretical kinetic modelling to identify the steps where these harmful intermediates arise. A similar approach has previously shown a direct causal link between amyloid-beta 42 oligomer formation and its pathological effect^52^. Our results show that αSyn’s damaging ability arises predominantly from αSyn oligomeric intermediates generated via secondary nucleation, rather than from mature fibrils. Pre-formed αSyn oligomer containing aggregation mixture did not influence this process; only αSyn fibrillar seeds were capable of amplifying the generation of toxic oligomers, which underlies their pathogenic potential. This may reflect the inherent instability of αSyn oligomers, which tend to dissociate into monomers^53^, in contrast to the structural stability of fibrils. Thus, sustained toxicity requires the continual generation of αSyn oligomers, a process that only fibrillar aggregates can effectively drive via secondary nucleation.

At first glance, these results may appear to challenge the prevailing view that αSyn oligomers are the principal toxic species. However, rather than contradicting this model, our results support and refine it. Previous studies directly comparing αSyn oligomers and fibrils, have relied on isolated systems^40^ or short-term assays^39^ focused on the immediate effects of exogenously added species. For instance, short-term studies (e.g., 20 min) in SH-SY5Y cells showed that exogenous oligomers are acutely more damaging to membranes than fibrils^39^. Yet longer-term experiments (>24 hours) in the same models reveal greater accumulation of endogenous aggregates in response to fibrillar seeds^54^. These observations suggest that αSyn oligomers may indeed be more harmful in settings requiring rapid, direct interactions, such as plasma membrane disruption^40^, where sustained amplification is not required. In contrast, the progressive cellular dysfunction observed in PD such as mitochondrial impairment and LB-like pathology, relies on the continual generation of toxic oligomers. Fibrillar seeds uniquely support this process through secondary nucleation, acting as catalytic scaffolds that continually replenish toxic oligomers. This distinction reconciles our data with previous findings, dissolves long-standing dichotomies between oligomer and fibril toxicity and offers a mechanistic explanation for how fibrillar seeds drive progressive αSyn pathology (**Figure 7A**).

**Figure 7.**
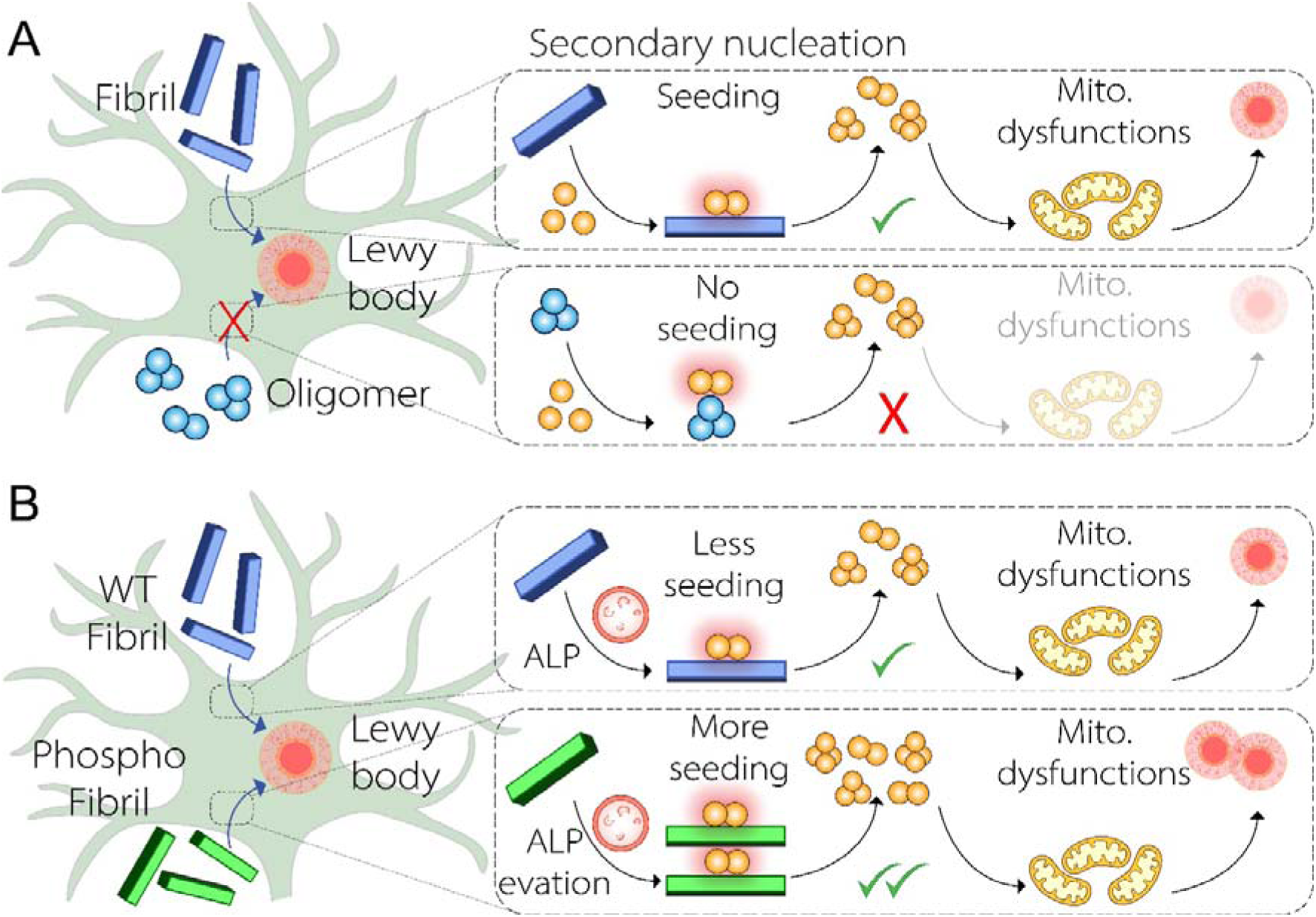
**(A)** Schematic showing how fibrillar αSyn seeds catalyze secondary nucleation, generating toxic oligomers and driving mitochondrial dysfunctions and LBs formation. In contrast, samples collected at the aggregation time point corresponding to peak oligomer levels fail to trigger this amplification cascade. **(B)** Phosphorylated αSyn fibrils evade autophagy-lysosomal degradation, persist longer, and induce greater seeding compared to WT fibrils, contributing to enhanced mitochondrial dysfunctions and LB-like pathology.

Another key finding of our study is that phosphorylated αSyn fibril seeds induce greater dysfunctions than their WT counterparts, despite showing comparable seeding efficiency. This finding is consistent with earlier reports showing that pre-formed WT αSyn fibrils induce less robust LB-like pathology than phosphorylated pS129 fibrils^55^ or those isolated from human PD brain^56,57^. We traced this differential susceptibility to altered engagement with cellular degradation pathways. Under basal conditions, αSyn aggregates are cleared via both the ALP and the UPP, consistent with previous studies^58–60^. However, during αSyn overload, we found partial impairment of both ALP and UPP, consistent with prior studies^59,61,62^. While both WT and phosphorylated fibrils were degraded via the UPP, we found that WT fibrils are efficiently trafficked to lysosomes and degraded via ALP. In contrast, phosphorylated fibrils exhibit reduced sensitivity to ALP-mediated degradation, similar to previous reports^60^. Since ALP becomes the dominant clearance route during αSyn overload^61,63^, similar to seeding conditions, this reduced susceptibility allows phosphorylated fibrils to persist longer inside cells. Their prolonged intracellular persistence, relative to WT fibrils, leads to greater overall production of toxic oligomers via secondary nucleation, driving sustained organelle damage and the formation of LB-like species formation. Over time, this persistent proteostatic burden may overwhelm the cell’s degradation capacity, triggering compensatory clearance routes such as extracellular release^25,54^. This, in turn, expands the pool of seeding-competent species extracellularly and reinforces a self-propagating cycle of pathology spread in the PD brain **(Figure 7B**).

An additional contributor to the heightened toxicity of phosphorylated αSyn fibrils may be the increased membrane disrupting potential of the new oligomer they generate. Phosphorylated fibril seeds produced more phosphorylated αSyn aggregates in iNLs (**Figure S6A-B**). In parallel, phosphorylated αSyn oligomers caused significantly greater membrane disruption than WT oligomers in cardiolipin-containing vesicles, which is specific to mitochondrial membranes and contribute to αSyn aggregate induced membrane damage^48,49,64^. Therefore, the observed increase in mitochondrial dysfunction (**Figure 3F,H**) and elevated colocalization of αSyn with TOMM20 (**Figure 3K**) in phosphorylated fibril-seeded iNLs is likely driven in part by the enhanced mitochondrial membrane-disruptive capacity of the aggregates formed from phosphorylated seeds.

While we showed that phosphorylated αSyn are more potent seeds than its WT counterpart due to higher persistence, our data indicate that total phosphorylated αSyn burden alone does not predict cellular dysfunctions. Absolute levels of phosphorylated αSyn did not distinguish diseased from control lines (**Figure 2B**), nor were they sufficient to induce mitochondrial dysfunctions and LB-like pathology, even when cells were seeded with aggregation time point peaked with phosphorylated αSyn oligomers (**Figure 3F-K**). These results suggest that pS129 is not inherently pathogenic, but that its detrimental effects emerge only when the balance between soluble and aggregated αSyn is disrupted. This interpretation is in agreement with recent studies proposing physiological roles for pS129 αSyn under normal conditions, including the regulation of synaptic function and αSyn turnover^17,18^. Moreover, although we aimed to generate pS129 αSyn using PLK3, the kinase also induced unintended phosphorylation at additional sites, suggesting that the observed effects may reflect the combined impact of multisite phosphorylation. Future studies are needed to disentangle the individual contributions of each modification to both normal αSyn function and pathological outcomes.

Extending our framework to the postmortem tissue, we found that total αSyn abundance did not correlate with LB burden in PD brains, mirroring our observations in iNL models. Instead, regions with more extensive LB pathology consistently exhibited higher seeding capacity. Given that only fibrillar αSyn species acted as potent seeds, these findings suggest that fibrillar seeds are likely harboured within LBs or closely associated with them. Kinetic modelling of seeded aggregation further revealed that estimated seed concentrations strongly correlated with regional LB burden and the spatiotemporal pattern of disease progression. While a recent study has linked seed amplification assay parameters such as the lag phase and number of positive replicates, to semi-quantitative measures of LB pathology^65^, our work directly links pathogenic seed concentrations to histologically defined LB burden. Together, these findings provide a molecular basis for the diagnostic power of αSyn seed amplification assays and establish a direct link between seeding capacity and disease trajectory in PD.

Collectively, our findings indicate that in sporadic PD, pathology is not simply dictated by αSyn expression levels or total aggregate load, but rather by a balance between physiological αSyn and its pathological conversion. This balance can be tipped by the introduction of exogenous seeds^66^, which trigger secondary nucleation and drive the continual formation of toxic oligomers, and thereby causing mitochondrial dysfunction, Lewy body formation, and eventual neurodegeneration. The potency and persistence of these seeds dictate the severity of the pathological cascade. Thus, what renders αSyn pathological is not its abundance per se, but its capacity to initiate self-amplifying toxic conversions. In sporadic PD, such seeds may arise from increased αSyn aggregation, impaired clearance, or transient cellular insults, such as inflammation^67^ or environmental stressors^68^, highlighting the urgent need to better understand these early initiating events.

In summary, our findings reveal that αSyn-induced dysfunction in PD is not a consequence of the static presence of oligomers or fibrils alone, rather emerges from their dynamic self-perpetuating interplay. Fibrillar αSyn acts as a catalytic scaffold, that accelerates the formation of neurotoxic oligomeric intermediate through secondary nucleation, a process that drives persistent LB pathology and organelle dysfunction. Our work reframes αSyn-induced damage in PD as a kinetic cascade, governed by the rate, persistence, and amplification of pathological species, rather than their absolute abundance. By mapping the molecular and temporal trajectory of αSyn seeded aggregation across cellular and anatomical contexts, our study offers a mechanistic foundation for targeting the most pathogenic steps of the PD cascade, with implications for both biomarker development and disease-modifying strategies.

### Limitations of the study

While this study advances understanding of how αSyn aggregation drives mitochondrial dysfunctions and Lewy pathology, several limitations remain. First, our kinetic modelling estimates total seed abundance in post-mortem tissue assuming either WT or phosphorylated αSyn, but both species coexist in the PD brain. We cannot resolve their individual contributions to seeding or how their relative abundance influences PD progression. Furthermore, we did not assess the contribution of C-terminally truncated αSyn which has shown to play role in LB pathology. Second, we lacked access to matched bio-fluids and post-mortem brain tissue, preventing direct comparisons between brain-resident seeds and diagnostics like SAAs. Third, this study was conducted using two sporadic PD lines, one SNCA triplication line, one PRKN mutant line, and four matched controls for each disease line. While this provides a unified mechanistic framework for different PD subtypes, inclusion of additional lines would further strengthen the generalizability of the findings. Finally, while our framework applies to sporadic and *SNCA*-linked PD, it may not fully capture mechanisms in genetic subtypes like *PRKN* or *LRRK2*, where αSyn pathology is variable. Extending this approach to broader genotypes will be important for understanding the role of αSyn across PD subtypes.

## Methods

### Fibroblast to iNPC culture

Fibroblast lines used in this study, derived from individuals diagnosed with Parkinson’s disease (PD) as well as neurologically healthy control (NHC) subjects, were obtained either via the Rutgers cell repository, Coriell cell repository or through the collection of forearm skin biopsies performed at the University of Sheffield (**Table 1**). All biopsies were obtained with appropriate ethical approval (Sheffield sourced cells are covered under the National Health Service National Research Ethics service (12/YH0367), and Rutgers cell repository and Coriell cell repository sourced cells are covered by MTA). Fibroblasts were cultured in Minimum Essential Medium (MEM) (Corning) supplemented with 10% fetal bovine serum (FBS) (Labtech), 100 U/mL penicillin (Lonza), 100 µg/mL streptomycin (Lonza), 50 µg/mL uridine (Millipore Sigma), 1 mM sodium pyruvate (Millipore Sigma), 0.1 mM non-essential amino acids (Fisher Scientific), and 0.1× MEM vitamins (Corning), and maintained at 37°C in a 5% CO₂ atmosphere. These fibroblasts were then directly reprogrammed into induced neuronal progenitor cells (iNPCs) using previously established protocols prior to this study^31,32,69^.

**Table 1.** Cell line information.

| Cell line | Mutation | Sex | Age at biopsy | Source |
| --- | --- | --- | --- | --- |
| **SNCA** | Triplication | F | 55 | Rutgers cell repository |
| **sPD1** | n/a | F | 56 | Sheffield |
| **sPD2** | n/a | F | 51 | Sheffield |
| **PRKN** | ARG42PRO;<br>EX3DEL / EX3DEL | M | 54 | Rutgers cell repository |
| **Control 1** | n/a | F | 55 | Rutgers cell repository |
| **Control 2** | n/a | M | 58 | Sheffield |
| **Control 3** | n/a | M | 55 | Sheffield |
| **Control 4** | n/a | M | 58 | Coriell cell repository |

### Differentiation of iNLs from iNPCs

iNPCs were cultured in Dulbecco’s modified essential media/ Ham’s F12 mixture Glutamax (DMEM/F12 Glutamax) (Fisher Scientific), supplemented with 1% N2 (Fisher Scientific), 1% B27 (Fisher Scientific),1% penicillin streptomycin (Lonza), and 40 ng/ml basic fibroblast growth factor (FGFb) (Peprotech). iNPCs were differentiated in iNLs as described in Schwartzentruber et al., 2020, Carling et al., 2020. At the end of the differentiation cells were assayed. The differentiation concentrations and timelines have been adjusted for some of the cell lines used in this paper to avoid cell detachment but remained consistent between matched pairs (**Table 2**)

**Table 2.** iNL differentiation timelines.

| Cell line | DAPT | FGF8/SAG | TGFβ/dcAMP/BDNF/GDNF |
| --- | --- | --- | --- |
| **SNCA** | 1 μM; 2 days | 37.5 ng/mL FGF8; 0.5 μM SAG; 5 days | 25 ng/mL BDNF/GDNF; 33 ng/mL TGFβ; 125 μM dcAMP; 7 days |
| **sPD1** | 1 μM; 2 days | 37.5 ng/mL FGF8; 0.5 μM SAG; 4 days | 25 ng/mL BDNF/GDNF; 33 ng/mL TGFβ; 125 μM dcAMP; 15 days |
| **sPD2** | 1 μM; 2 days | 37.5 ng/mL FGF8; 0.5 μM SAG; 10 days | 25 ng/mL BDNF/GDNF; 33 ng/mL TGFβ; 125 μM dcAMP; 5-8 days |
| **PRKN** | 2.5 μM; 2 days | 75 ng/mL FGF8; 1 μM SAG; 10 days | 25 ng/mL BDNF/GDNF; 33 ng/mL TGFβ; 125 μM dcAMP; 15 days |
| **Control 1** | 1 μM; 2 days | 37.5 ng/mL FGF8; 0.5 μM SAG; 5 days | 25 ng/mL BDNF/GDNF; 33 ng/mL TGFβ; 125 μM dcAMP; 7 days |
| **Control 2** | 1 μM; 2 days | 37.5 ng/mL FGF8; 0.5 μM SAG; 4 days | 25 ng/mL BDNF/GDNF; 33 ng/mL TGFβ; 125 μM dcAMP; 10 days |
| **Control 3** | 1 μM; 2 days | 37.5 ng/mL FGF8; 0.5 μM SAG; 10 days | 25 ng/mL BDNF/GDNF; 33 ng/mL TGFβ; 125 μM dcAMP; 5-8 days |
| **Control 4** | 2.5 μM; 2 days | 75 ng/mL FGF8; 1 μM SAG; 10 days | 25 ng/mL BDNF/GDNF; 33 ng/mL TGFβ; 125 μM dcAMP; 15 days |

### Immunocytochemistry

At the end of differentiation, iNPCs or iNLs were washed once with 1× PBS and fixed with 3.7% paraformaldehyde for 15 minutes at room temperature. Cells were then washed three times with 1× PBS and permeabilized with 0.2% Triton X-100 in PBS for 10 minutes. After permeabilization, cells were washed three times to remove Triton X-100 and blocked for 1 hour in 3% horse serum prepared in 1× PBS containing 0.1% Triton X-100. For iNPCs, cells were incubated overnight at 4 °C with primary antibodies against Pax6 (Abcam ab18102) and Nestin (Abcam ab5790) both diluted 1:200. For iNLs, primary antibodies against TuJ1 (Abcam ab190575, diluted 1:1000), DAT (Invitrogen PA5-78382, diluted 1:500), and TH (Biolegend 818006, diluted 1:200) were used. For experiments involving mitochondrial and α-synuclein detection in iNLs, primary antibodies (e.g., TOMM20; BD Biosciences 612278 diluted 1:1000, αSyn; Abcam Sigma AB5038 diluted 1:1000) were also incubated overnight at 4 °C. The following day, cells were washed three times with PBS and incubated with secondary antibodies (Invitrogen, diluted 1:1000 in blocking buffer) for 1 hour at room temperature. The secondary antibodies conjugated to Alexa Fluor 488, 568 or 647 and were applied for 1 hour, followed by 10 μM Hoechst 33258 (Sigma) for 10 minutes. After a final three PBS washes, cells were imaged.

### Imaging of and live and fixed cells

Imaging of both fixed and live cells cultured in black 96-well flat bottom plates was performed using the Opera Phenix High-Content Screening System (Revvity) equipped with a 40× water immersion objective. For live-cell imaging, environmental conditions were maintained at 37 °C with 5% CO₂. Image acquisition parameters were adapted based on the fluorophores used in each assay, with excitation lasers at 405 nm, 488 nm, 568 nm, and 647 nm. For each well, a minimum of 9 fields of view were acquired, each comprising 5-6 Z-planes, capturing a minimum of 100 cells per well. At least three wells were imaged per condition in each independent experiment.

### Mitochondrial phenotyping

For mitochondrial function assays, media was removed from live cells. Cells were incubated with a dye mixture in clear MEM without phenol red (Fisher Scientific), including TMRM 80 nM (Sigma), MitoTracker^TM^ Green 1µM (Invitrogen), LysoTracker^TM^ Deep Red 50 nM (Invitrogen) and Hoechst 33258 40µM. Cells were incubated with dyes for 1 hour at 37°C. Subsequently, dyes were removed, and clear MEM was added to the cells prior to imaging with the Opera Phenix High Content Screening System using the 40x water objective, at 37°C and 5% CO_2_.

### Blocking autophagy and proteasomal degradation

At the end of differentiation, iNLs were treated for 24 hours with either 10 µM Chloroquine (Millipore Sigma) to inhibit autophagy or 10 µM MG132 (Millipore Sigma) to inhibit proteasomal degradation. After 24 hours, media was replaced, and αSyn aggregates were added at a final concentration of 1 µM for 8 hours. Then media was collected, cells were washed with PBS and lysed.

### Analysis of cell imaging data

All images acquired on the Opera Phenix system were analysed using Harmony 5.2® High Content Analysis Software (Revvity). Analysis pipelines were customized to suit each experimental condition. Z-stacks were processed using maximum intensity projection, and flatfield correction was applied to minimize illumination artifacts. Nuclei were identified based on size, intensity, and morphology to exclude debris and non-viable cells. Cytoplasmic regions were segmented using either cytoplasm detection algorithms or whole-image region analysis, depending on the staining pattern. To quantify the percentage of cells positive for iNPC or iDNL markers, cytoplasmic fluorescence intensity was measured. A threshold was set based on background staining from secondary antibody-only controls, and cells exceeding this threshold were classified as marker-positive.

For live-cell imaging of mitochondrial and lysosomal compartments, mitochondria and lysosomes were segmented using MitoTracker™ Green, TMRM, and LysoTracker™ Deep Red signals. Segmented structures were filtered by size and intensity to exclude background and artifacts. Organelle counts were quantified per cell. Mitophagy index was evaluated by quantifying the colocalization of MitoTracker™ Green positive mitochondria and LysoTracker™ Deep Red positive lysosomes, as a percentage of total mitochondria. This index was used as a surrogate for mitochondrial degradation via lysosomes. This imaging-based approach to assess mitochondrial function and mitophagy has been validated in prior studies^31,32,69^.

### Cell lysis

Cell media was collected, and cells were washed with 1x PBS. Then ice-cold lysis buffer (20 mM Tris-HCl pH 7.4, 150 mM NaCl, 1% TritonX-100, 10% glycerol, supplemented with 1 mM EDTA, 1 mM PMSF, 1 mM PIC) was added and cells were scraped and collected. Samples were needled 10 times using 25-gauge needles and incubated on ice for 10 minutes. Cell lysates were then centrifuged at 14,000 g for 5 minutes at 4°C. Supernatant was collected and total protein concentration was determined.

### Total protein measurement

Total protein concentration of cell lysates was determined using the Pierce™ Bicinchoninic acid (BCA) Protein Assay Kit (Thermo Scientific), following the manufacturer’s instructions. Bovine serum albumin (BSA) standards were prepared in PBS to generate a standard curve. Cell lysate samples were diluted 1:10 in PBS to ensure concentrations fell within the linear range of the assay. For each well of a clear 96-well microplate, 25 µL of standard or diluted sample was added in triplicate, followed by 200 µL of working reagent (prepared by mixing BCA Reagent A and B at a 50:1 ratio). The plate was briefly shaken for 30 seconds to ensure proper mixing and then incubated at 37 °C for 30 minutes. After incubation, the plate was cooled to room temperature, and absorbance was measured at 562 nm using a Clariostar Plus microplate reader (BMG Labtech). Protein concentrations of the samples were calculated based on the BSA standard curve.

### Meso-scale discovery assay

MSD assay was used to quantify different forms of αSyn, including total, aggregated, and phosphorylated species. For capture, 4 µg/mL of Syn211 (Abcam, ab138501), which recognizes both monomeric and aggregated αSyn, was used for total and aggregated αSyn detection. For phosphorylated αSyn at serine 129 (pS129), 3Lµg/mL of anti-αSyn pS129 antibody (EP1536Y, Abcam, ab209422) was used. Capture antibodies were coated onto MSD plates and incubated overnight at 4 °C in PBS without agitation. Following incubation, plates were washed three times with 200 µL of PBST (0.1% Tween-20 in PBS) and then blocked with 1% BSA in 1% PBST for 2 hours at 37 °C with shaking. After an additional three washes, 25 µL of sample was added per well and incubated for 2 hours at room temperature with orbital shaking at 800 rpm. Subsequently, plates were washed again and 40 µL of detection antibody (2 µg/mL) was added. For total and pS129 αSyn, anti-human αSyn detection antibodies were used. For aggregated αSyn, Syn211 was used for both capture and detection; the use of identical monoclonal antibodies in this format selectively detects aggregate forms due to the requirement for multivalent binding. After a final set of washes, 200 µL of MSD Read Buffer was added to each well. Plates were read using a MESO QuickPlex SQ 120 multiplex imager. As signal intensity can be affected by the sample matrix (e.g., cell or tissue lysates), all measurements were performed in a semi-quantitative manner. Results are normalized to total protein concentration determined by the BCA assay.

### Single Molecule Pulldown assay (SiMPull) assay

SiMPull prepared coverslips were placed into a humid chamber. For biotin-neutravidin linkage, 0.2 mg/ml neutravidin was diluted in PBST and added to each well for 5 minutes. The wells were then washed twice with PBST and once with 1% Tween in 1x PBS. The biotin capture antibody (Biotinylated Syn211) was immobilised onto the coverslip surface through biotin-neutravidin linkage, the antibody was diluted to 10 nM in PBS-BSA and incubated for 10 minutes. Each well was then washed twice with PBST and once with 1% Tween in PBS. Samples (diluted cell lysate) were then applied to the wells and incubated in the humid chamber for at least 1 hour. After sample incubation, each well was washed twice with PBST and once with 1% Tween in 1x PBS. Imaging antibodies conjugated to dye (Alexa-Fluor 637 labelled Anti-αSyn aggregate antibody, MJFR-14-6-4-2, Alexa-Fluor 561 labelled MJFR phosphorylated αSyn specific antibody, LB marker Alexa-Fluor 488 conjugated) were then applied onto the wells at a concentration of 5 nM (αSyn antibodies) or 10 nM (LB or organelle protein marker antibodies) and incubated for 30 minutes protected from light. Each well was then washed twice with PBST, before a second gasket was placed onto the top of the coverslip, and 1x PBS applied into the wells. The coverslip was sealed with a second argon cleaned coverslip and imaged using a Nikon Ti2 TIRF microscope.

### Antibody-dye conjugation for SiMPull assay

Unlabelled antibodies were conjugated for use in the SiMPull assay using Lightning-Link® conjugation kits (Abcam), following the manufacturer’s instructions. Antibodies were labelled with Biotin, AF488, AF568, or AF647, depending on the application. Briefly, 1 µL of modifier reagent was added to 10 µL of unlabelled antibody and mixed gently. This antibody–modifier solution was then transferred to the lyophilized conjugation mix and gently pipette-mixed to dissolve the contents. The reaction mixture was incubated overnight in the dark at room temperature to allow conjugation. Following incubation, 1 µL of quencher reagent was added to deactivate any unbound dye or reactive species, and the solution was gently mixed again. After a 5-minute final incubation, the labelled antibodies were ready for use and stored at 4 °C. According to the manufacturer, labelled antibodies remain stable under these conditions for up to 18 months.

### Total internal reflection fluorescence (TIRF) microscopy

Single-aggregate imaging and SiMPull assays were performed using a custom-built TIRF microscope based on the Nikon Ti2 Eclipse platform. Excitation was achieved using laser lines at 405 nm, 488 nm, 561 nm, and 638 nm. Each laser passed through a neutral density filter to control intensity, was expanded using beam-expanding optics, and circularly polarised via a quarter-wave plate. A beam shaper (Asphericon) converted the Gaussian beam into a top-hat profile for uniform field illumination. The expanded beams were directed to the back focal plane of a 100×, NA 1.49 oil-immersion TIRF objective. Emitted fluorescence was collected through the same objective, separated from excitation light using a beam splitter, passed through appropriate bandpass filters, and detected by a Prime 95B sCMOS camera. Field-of-view selection and image acquisition were controlled by a custom automated stage-scanning program to eliminate user bias. This program also enabled precise re-positioning for multi-channel colocalization measurements. All microscope functions and image capture were managed using Micro-Manager open-source software. For each biological replicate, a minimum of nine fields of view were acquired. Each field was imaged for 50 frames with an exposure time of 50 ms per frame.

### Analysis of colocalization data

Averaged images obtained using multiple excitation channels were processed in Fiji using the ComDet3 plugin. Spots from two or three channels were classified as colocalized if the distance between their Gaussian-fitted centres was ≤2 pixels. Colocalization by aggregate count was calculated as the proportion of colocalized spots relative to the total spot count in the given channel.

### Recombinant α-Synuclein expression and purification

Recombinant human αSyn was expressed and purified following a modified version of established protocols. Briefly, αSyn was expressed from the pT7-7 plasmid in *E. coli* BL21 (DE3) cells grown in 6LL 2×YT medium with 100 μg/mL ampicillin. Protein expression was induced with IPTG, and cultures were incubated overnight at 28 °C. Cells were harvested by centrifugation (6,240L×Lg), resuspended in lysis buffer (10 mM Tris pH 8.0, 1 mM EDTA, 1 mM PMSF), lysed by sonication, and boiled for 20Lminutes at 85 °C. After centrifugation (39,000L×Lg), the supernatant was treated with streptomycin sulfate (10 mg/mL) for 15Lminutes at 4 °C and centrifuged again. Ammonium sulfate (0.36 g/mL) was added to the supernatant, stirred for 30Lminutes at 4 °C, and centrifuged once more. The resulting pellet was resuspended in 25LmM Tris (pH 7.7), dialyzed overnight, and purified by ion-exchange chromatography (Q Sepharose HP column) using 25 mM Tris buffers with or without 1.5 M NaCl. Fractions containing αSyn were subjected to size exclusion chromatography (HiLoad 26/600 Superdex 75 pg). Protein concentration was measured spectrophotometrically (ε280 = 5,600LM⁻¹Lcm⁻¹). The N122C mutant was purified similarly with 3 mM DTT in all buffers. For experiments requiring fluorescently labelled protein, Alexa Fluor 488–conjugated αSyn was purchased from a commercial vendor (AnaSpec).

### Phosphorylation protocol

Phosphorylation of α-synuclein (αSyn) at serine 129 (S129) was performed following a previously published protocol^36^. Briefly, a phosphorylation buffer was freshly prepared containing 50 mM HEPES, 1 mM MgCl₂, 1 mM EGTA, and 1LmM DTT. The reaction mixture consisted of 230 µM αSyn, 2 mM Mg-ATP, and 1 µL of PLK3 kinase, added to the phosphorylation buffer. The solution was mixed thoroughly by pipetting and incubated at 30 °C for 12 hours without agitation. Successful phosphorylation at S129 (P-αSyn) was confirmed by western blotting using the MJF-R13 phospho-specific α-synuclein antibody (Abcam, ab168381) and mass spectrometry.

### Western blot

To confirm phosphorylated of αSyn, 2 µg of phosphorylated and non-phosphorylated αSyn was mixed with Laemmli buffer and boiled at 95°C for 5 minutes to denature the proteins. Samples were loaded onto a 4-20% Mini-PROTEAN polyacrylamide gel (BioRad), alongside a protein ladder. Proteins on the gel were then transferred to a nitrocellulose membrane using a Criterion blotter (BioRad) in transfer buffer (50 mM Tris HCl, 20% methanol). The transfer was ran at 100 V for 30 minutes. After the transfer, the membrane was fixed with 2% formaldehyde for 30 minutes at room temperature. The membrane was then blocked in 5% milk in TBST for 1 hour at room temperature, before incubating primary antibodies (Syn211; Abcam ab206675 diluted 1:1000, MJFR1 Abcam ab209420 diluted 1:1000, MJFR-Phospho Abcam ab168381) in blocking buffer at 4°C overnight. The next day, the membrane was washed three times in TBST, with each wash lasting 10 minutes. Next, fluorescent secondary antibodies (Jackson ImmunoResearch, Anti-mouse Alexa-Fluor 790, Anti-rabbit Alexa-Fluor 680) were diluted 1:50,000 in TBST and incubated on the membranes for 1 hour at room temperature protected from light. Subsequently, the membranes were washed three times in TBST before imaging using the LI-COR Odyssey Fc imaging system.

### Mass spectrometry analysis of α-synuclein phosphorylation

Protein from in vitro phosphorylation assays of αSyn +/- PLK3 was added to equal volumes of 2X S-trap lysis buffer (10% SDS and 100 mM Triethylammonium bicarbonate (TEAB) and reduced by the addition of Tris(2-carboxyethyl) phosphine hydrochloride (TCEP) to a final concentration of 5 mM and samples were heated to 70°C for 15 minutes. Iodoacetamide (IAA) was then added to a final concentration of 10 mM and samples were incubated at 37°C in the dark for 30 minutes. 2.5μL of 12% phosphoric acid and 165μL of S-Trap binding buffer (90% methanol, 100 mM TEAB pH 7.1) was then added to each sample. Samples were then loaded into S-Trap columns (ProtiFi) 150 μL at a time by centrifugation at 4000 rpm for 30 seconds. Samples were washed four times with 150μL of S-Trap binding buffer through centrifugation at 4000rpm for 30 seconds. Trypsin (Pierce, sequencing grade) was added to each sample at a 1:10 ratio and digestion was allowed to proceed at 47°C for 1h followed by 37°C for 1 hour. Peptides were eluted with the addition of 40μL of 50mM TEAB, 40μL of 0.2% aqueous formic acid and then 40μL of 50% acetonitrile with 0.2% formic acid followed by centrifugation at 4000 rpm for 30 seconds for each elution. Pooled eluted peptides were dried in a vacuum concentrator and resuspended in 0.5% formic acid for LC-MS/MS analysis. To generate suitable peptides to detect phosphorylation of α-synuclein at Ser129, ProAlanase (Promega) was added to the tryptically digested peptides and digested at 37°C for 1.5 hours. Each sample was analysed using nanoflow LC-MS/MS using an Orbitrap Exploris 480 (Thermo Fisher) mass spectrometer with an EASY-Spray source coupled to a Vanquish LC System (Thermo Fisher). Peptides were desalted online using a PepMap Neo C18 nano trap column, 300μm I.D.X 5mm (Thermo Fisher) and then separated using an EASY-Spray column, 50cm X 75μm ID, PepMap Neo C18, 2μm particles, 10Å pore size (Thermo Fisher). The Orbitrap Exploris 480 was operated in positive mode with a DDA cycle time of 2s. MS1 spectra were acquired at a resolution of 60,000 at m/z 200 with a standard AGC target. The most abundant multiply charged ions (2-6+) in a given chromatographic window subjected to HCD fragmentation with a collision energy of 30% and dynamic exclusion set to automatic. The MS2 AGC target was set to standard, and MS2 spectra were measured with a resolution setting of 15,000 at m/z 200 a standard AGC target. To improve detection of the c-terminal peptide of αSyn bearing phosphorylation of Ser129, charge states 1-6 were subjected to fragmentation.

Raw mass spectrometry data files were processed with MaxQuant version 1.6.10.43. Data were searched against an Uniprot E coli (March 2021) sequence database with the sequences of human *SNCA* and PLK3 using the following search parameters: digestion set to Trypsin/P with a maximum of 2 missed cleavages, methionine oxidation, N-terminal protein acetylation and Phospho (STY) as variable modifications, cysteine carbamidomethylation as a fixed modification. A first search precursor tolerance of 20 ppm and a main search precursor tolerance of 4.5 ppm was used for FTMS scans and a 0.5 Da tolerance for ITMS scans. A protein FDR of 0.01 and a peptide FDR of 0.01 were used for identification level cut-offs. For samples subjected to a second digestion with ProAlanase, cleavage after alanine, proline and arginine and lysine were allowed. Class I phosphorylation sites were filtered to a localisation probability of >0.75 and a score difference of >5.

### Measurement of αSyn aggregation kinetics

Firstly, 1 mM ThT stock solution was prepared by dissolving the dye in absolute ethanol and subsequently diluted to 200 µM in 0.02 μm-filtered PBS. The exact concentration of the working solution was confirmed by measuring absorbance at 412 nm (ε = 36,000 M⁻¹cm⁻¹) using a Nanodrop spectrophotometer. Before running the aggregation, WT or phosphorylated αSyn monomer was ultracentrifuged at 20,000 × g for 1 hour at 4 °C to remove pre-existing aggregates. Aggregation reactions were conducted in 300 µL volumes containing 25 mM Tris-HCl (pH 7.4), 100 mM NaCl, and 0.01% NaN_3_, incubated at 37 °C with continuous shaking at 200 rpm. Aliquots (10 µL) were collected at defined time points and flash-frozen for subsequent analysis. ThT fluorescence measurements were performed simultaneously for all time points using a Clariostar plate reader to reconstruct aggregation kinetics. For seeded experiments, preformed αSyn fibrils were sonicated for 10 minutes to generate short fibril fragments, which were then added to fresh monomer to initiate seeded aggregation. For dye-labelled αSyn aggregates preparation, AF488 conjugated αSyn was mixed at a 1:9 ratio (10% labelled, 90% unlabelled) with unlabelled αSyn. Aggregation was performed in the same conditions, and time-matched aliquots were collected for aggregate isolation.

### Single-aggregate imaging using ThT

For characterization the αSyn aggregates (WT or phosphorylated) were generated under identical conditions as described above, excluding ThT dye. At designated time points, aliquots were collected and diluted to 200 nM in filtered PBS for imaging. Glass coverslips were plasma-cleaned with argon for at least 30 minutes (Harrick Plasma), coated with 0.01% poly-L-lysine solution for 30 minutes, and washed three times with filtered PBS. Diluted aggregate samples were then incubated on the treated coverslips in PBS containing 5 µM ThT for 10 minutes to allow surface adsorption. After incubation, chambers were immediately mounted onto the microscope stage. Imaging was performed in TIRF mode using 405 nm laser excitation at a frame rate of 50 ms. Emitted fluorescence was collected through a 405 nm long-pass razor-edge filter. All images were processed and analysed using Fiji (ImageJ).

### Seeding in cells

At the end of differentiation, WT and phosphorylated αSyn in vitro formed aggregates taken at 36 hours (oligomers) and 7 days (fibrils-sonicated) were added to healthy control iNLs cell media at a final concentration of 1 µM monomer equivalent. Cells were incubated at 37°C with 5% CO_2_. After 24 hours, the cell media was aspirated and replaced with fresh media, and cells were placed back into the incubator. After a further 24 hours the cells were assayed.

### Data analysis of cellular uptake assay

Images were captured from randomly selected fields across three wells per condition and were analysed using Harmony High-Content Imaging software. Two channels were used: 561 nm to visualize cell boundaries via TMRM staining, and 488 nm to detect WT or phosphorylated αSyn labelled with AF488. Cell boundaries in iNLs were defined based on the TMRM, and the resulting masks were applied to the αSyn channel to quantify fluorescence intensity, enabling measurement of αSyn uptake.

### Membrane permeabilization assays

Membrane permeabilization was assessed using a previously established protocol^42,70^. Vesicles were prepared to mimic mitochondrial membrane composition, consisting of 20% (10% or 0%) 16:0–18:1 PC (Avanti, 850457), 40% 16:0–18:1 PE (Avanti, 850757), 20% 18:1 cardiolipin (Avanti, 710335), 3% 16:0 SM (Avanti, 860584), 3% 16:0–18:1 PI (Avanti, 850142), 3% 16:0-18:1 PS (Avanti, 840034), and 1% biotinylated 18:1–12:0 Biotin-PC (Avanti, 860563C). For 10% and 0% cardiolipin conditions, other lipids were adjusted proportionally. Vesicles were extruded through a 100 nm polycarbonate membrane (Avanti, 610005) following ten freeze-thaw cycles and hydrated in 100 μM Cal-520 dye (Stratech, 21141) in 50 mM HEPES buffer of pH 6.5. They were immobilized on coverslips (VWR, 6310122) cleaned via argon plasma (Deiner Zepto One) and coated with a 100:1 mixture of PLL-g-PEG (20 kDa PLL with 2 kDa PEG and 3.5 Lys/PEG chain, SuSoS) and PLL-g-PEG-biotin (20 kDa PLL with 2 kDa PEG and 50% 3.4 kDa PEG-biotin, SuSoS) at ∼1 mg/mL. Next, 50 µL of 0.1 mg/mL NeutrAvidin (ThermoFisher, 31000) in HEPES buffer was added to the coverslips, followed by vesicle immobilization. Imaging was performed first in Ca²⁺-containing buffer (ThermoFisher, #21083027) to establish baseline fluorescence (F_blank_), then after 15-minute sample exposure (F_sample_), and finally after ionomycin treatment (Cambridge Bioscience, 1565-5) to achieve Ca²⁺ saturation (F_ionomycin_). Relative Ca²⁺ influx was calculated as: (F_sample_ − F_blank_)/(F_ionomycin_ − F_blank_). Cal-520 fluorescence was filtered (BLP01-488R-25 and FF01-520/44-25, Laser 2000) and imaged using a Photometrics Prime 95B sCMOS camera. Acquisition was performed at ∼10 W/cm² power density with a scan speed of 20 Hz.

### Kinetic modelling of aggregation and oligomer formation

We derive general rate equations for filament assembly that proceed through an intermediate oligomeric species, explicitly accounting for oligomer dissociation. Secondary processes, such as secondary nucleation, are also included. Although only a single type of oligomer is considered here, assuming a uniform oligomer population often serves as a useful and insightful approximation. Free monomer concentration is *m*(*t*), fibril mass concentration is *M*(*t*), and fibril number concentration is *P*(*t*). The oligomer number concentration is *S*(*t*). The sources of oligomers are primary nucleation, with rate 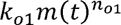, where *k_o_*_1_ is the rate constant of primary nucleation and *n_o_*_1_ is the reaction order of primary nucleation, and secondary nucleation, with rate 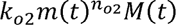, where *k_o_*_2_ is the rate constant of secondary nucleation and *n_o_*_2_ is the reaction order of secondary nucleation. We consider only secondary nucleation as the secondary process, since it contributes more significantly than fragmentation^45^. The sinks of oligomers are oligomer conversion to fibrils, with rate 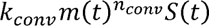, where *k_conv_* is the conversion rate constant and *n_conv_* is the conversion reaction order, and oligomer dissociation back to monomers, with rate *k_d_*_2_*M*(*t*)*S*(*t*), where *k_d_*_2_ is the rate constant of fibril-mediated dissociation. We consider only fibril-mediated dissociation here, as the direct dissociation rate is much slower^71^. Fibril number concentration *P*(*t*) mainly increases via oligomer conversion. Fibril mass concentration *M*(*t*) mainly increases via elongation, where *k*_+_ is the elongation rate constant. We ignore depolymerization, which is the reverse of elongation, as the monomer concentration at equilibrium is very low in the systems studied and therefore does not significantly affect the kinetics. We also disregard filament annealing, which is the reverse of fragmentation, as it has a negligible effect on the time profiles of the measured quantities *M*(*t*) and *S*(*t*). We assume that the mass concentration of oligomers is much smaller than the total protein concentration; therefore, the mass conservation law does not include the oligomer mass concentration. *m_tot_* is the total protein mass concentration in system.

The model can be described by the following moment equations:

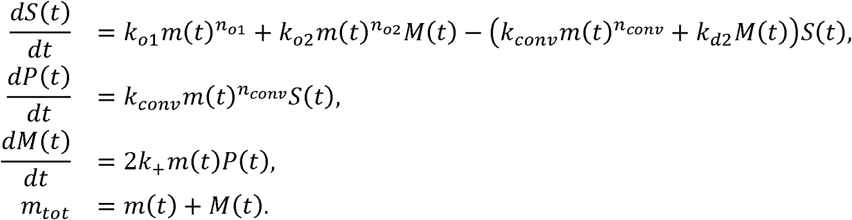

### Case selection

Post mortem human brain tissue was provided by the Sheffield Brain Tissue Bank (SBTB), University of Sheffield which has ethical permission to function as a Research Tissue Bank from the Wales Research Ethics Committee 5 (Reference 24/WA/0052). Informed consent was obtained from all donors in life or their relatives. Tissue samples were collected from five individuals with clinically and pathologically confirmed PD and five neurologically healthy control subjects (**Table 3**).

**Table 3.**
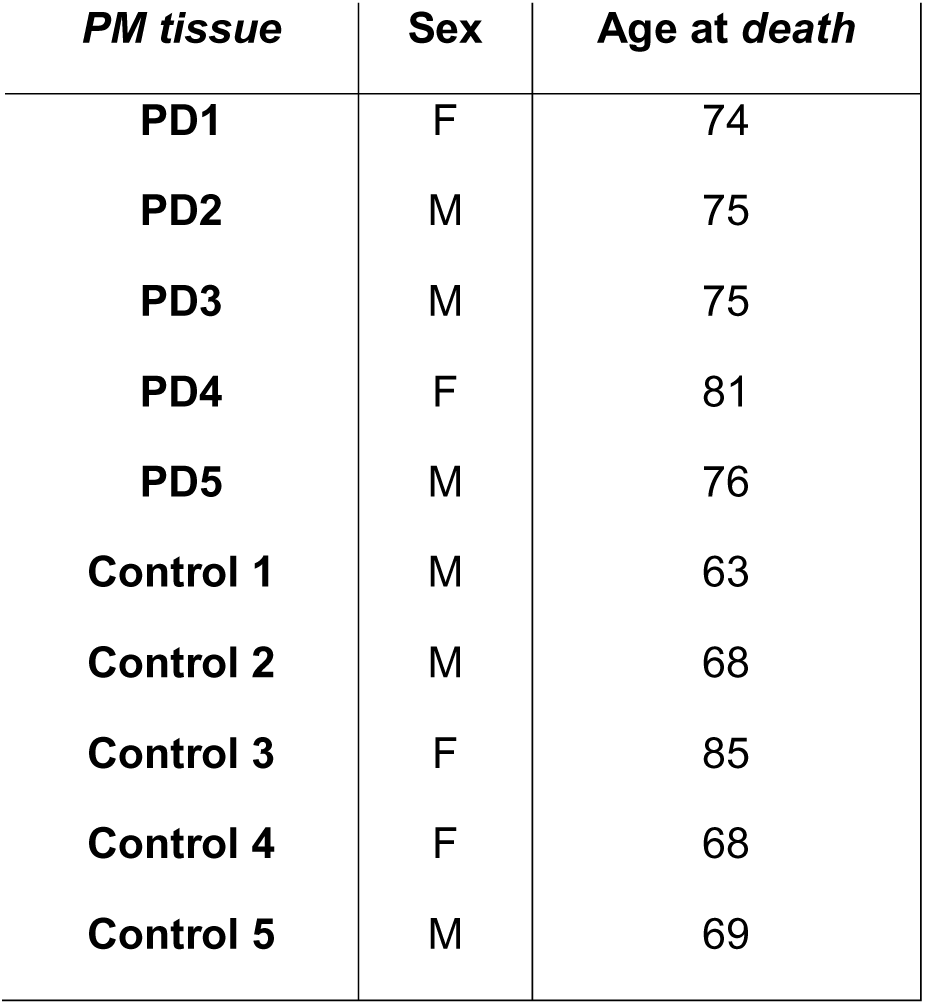
PM tissue information.

| *PM tissue* | *Sex* | *Age at death* |
| --- | --- | --- |
| **PD1** | F | 74 |
| **PD2** | M | 75 |
| **PD3** | M | 75 |
| **PD4** | F | 81 |
| **PD5** | M | 76 |
| **Control 1** | M | 63 |
| **Control 2** | M | 68 |
| **Control 3** | F | 85 |
| **Control 4** | F | 68 |
| **Control 5** | M | 69 |

Since we aimed to determine whether αSyn seeding activity correlates with LB burden, we strategically selected brain regions known to be affected at distinct stages of PD: the substantia nigra, cingulate gyrus, and neocortex. These regions represent sequential stages along the pathological trajectory, allowing us to capture stage-specific changes in both seeding activity and LB accumulation. For each region, we required two types of tissue from the same individuals: *(i)* formalin-fixed, paraffin-embedded (FFPE) tissue for immunohistochemical quantification of LBs and *(ii)* flash-frozen tissue for αSyn seeding assays. Formalin fixation can introduce chemical cross-links that alter αSyn aggregate conformation and seeding activity. Therefore, only flash-frozen samples preserve the native aggregation state required for functional assays. Conversely, FFPE sections are essential for reliable histological detection and quantification of LB pathology.

To enable both assays from anatomically matched regions within the same brains, each case was sagittally dissected. Half of the brain was formalin fixed, dissected and selected blocks processed to paraffin, while the contralateral hemibrain was coronally sliced and immediately flash-frozen between liquid nitrogen-cooled teflon-coated copper plates. For LB quantification, FFPE sections were obtained from the midbrain (for substantia nigra), cingulate gyrus, and neocortex. For αSyn seeding assays, flash-frozen tissue from the same cases was used.

### Immunohistochemistry of postmortem PD tissue

Five-micron-thick sections were cut from FFPE human tissue blocks from PD patients. Immunohistochemistry was performed using an antibody specific for phosphorylated α-synuclein (phosphorylated αSyn; BioLegend, 825701). Tissue sections were dewaxed in xylene and rehydrated through a graded ethanol series. Endogenous peroxidase activity was quenched by incubation in 3% hydrogen peroxide in methanol for 10 minutes. Antigen retrieval was performed using a pH 6.0 citrate buffer, with samples microwaved at 100% power for 10 minutes. This was followed by a 3-minute incubation in 100% formic acid to further expose antigenic sites. To minimize nonspecific binding, blocking was performed using 2.5% normal horse serum (VECTASTAIN® Elite ABC-HRP Kit, Mouse IgG, Vector Laboratories, Cat. No. PK-6102) for 30 minutes at room temperature. The primary phosphorylated αSyn antibody was diluted 1:500 and incubated with the sections overnight at 4 °C. The next day, sections were incubated with a biotinylated horse anti-mouse IgG secondary antibody for 30 minutes, followed by incubation with the avidin–biotin–peroxidase complex from the same kit for an additional 30 minutes. Signal was visualized using 3,3′-diaminobenzidine (DAB; Vector Laboratories, Cat. No. SK-4100) as the chromogen and hydrogen peroxide as the substrate. DAB reaction was allowed to proceed for 3 minutes. Sections were counterstained with hematoxylin, dehydrated through graded ethanol solutions, and mounted with dibutylphthalate polystyrene xylene (DPX). Whole-slide images were scanned using the Hamamatsu NanoZoomer XR slide scanner at 20× magnification. Cropped regions were prepared using QuPath, and scale bars were added in ImageJ. Manual quantification of Lewy bodies across the entire scanned sections was performed using Qu Path’s point annotation tool.

### Preparation of brain homogenates

Frozen brain tissue samples (∼200 mg) were homogenized using the Precellys Evolution tissue extractor with Precellys 2 mL Lysing Kits (CK14, optimized for soft tissue) in 400 µL of homogenization buffer supplemented with protease and phosphatase inhibitors (Halt™ Protease & Phosphatase Inhibitor Cocktail, Thermo Scientific, Cat. No. 78440). The generic soft tissue homogenization program was used: two cycles at 5500 rpm for 30Lseconds each, with a 30-second pause between cycles. Following homogenization, samples were placed on ice for 15 minutes and then centrifuged at 13,000 rpm for 20 minutes at 4 °C using a benchtop centrifuge. The resulting supernatant was collected, protein concentration was measured, and the homogenates were aliquoted and stored at −80 °C until further use. For αSyn seeding assays, aliquots were thawed on ice and diluted approximately 1:2 in assay buffer, with input normalized to the original tissue weight.

### Expression and Purification of Bri2 BRICHOS R221E Monomer

BRICHOS R221E was prepared following a previously published protocol^50^. In short, BRICHOS variant was generated by site-directed mutagenesis using the wild-type His_6_-NT*-Bri2 BRICHOS (113-231) plasmid as a template. The construct was expressed in Shuffle T7 *E. coli* cells grown in LB medium supplemented with kanamycin (15 ug/ml) at 30 °C. Protein expression was induced with 0.5 mM IPTG at an optical density of 0.8, followed by overnight incubation at 20L°C. The next day, the cells were harvested by centrifugation (5.000 x g, 20 min at 4 °C). After harvesting, cells were resuspended in 20 mM Tris buffer (pHL8.0), lysed by sonication, and clarified by centrifugation (24.000 x g, 4 °C for 30. min). The NT*-BRICHOS fusion protein was purified using Ni-NTA affinity chromatography. Thrombin digestion was performed during overnight dialysis to cleave off the His_6_-NT* solubility tag leaving an additional glycine and serine at the N-terminal part of BRICHOS. The His_6_-NT* protein was removed by a second Ni-NTA purification step. The flow through containing purified BRICHOS was heat treated for 10 min at 60 °C and monomeric protein was isolated by size-exclusion chromatography (Hiload Superdex 75 PG). The final purified protein was concentrated to the desired concentration and stored at −70 °C until further use.

### Statistical Analysis

All statistical analyses were performed using Origin 9.0 (OriginLab). Data are presented as mean values, with error bars indicating the standard deviation from biological replicates. For comparisons between two groups, unpaired two-tailed Student’s *t*-tests were used. For comparisons involving more than two groups, one-way ANOVA followed by Tukey’s post hoc test was performed. Statistical significance was defined as follows: *p* < 0.05 = *, *p* < 0.01 = **, *p* < 0.001 = ***, and *p* < 0.0001 = ****.

### Lead contact

Further information and requests for resources and reagents should be directed to and will be fulfilled by the lead contact, Dr. Suman De.

## Data and availability

Data and analysis code reported in this paper will be shared by the lead contact upon request. Any additional information required to reanalyse the data reported in this paper is available from the lead contact upon request.

## Material availability

This study did not generate new, unique reagents.

## Supporting information

Mass spec data

## Acknowledgments

We would like to thank the participants from which the original skin biopsies were taken. We are also grateful to the Sheffield Brain Tissue Bank for supplying the tissue and to those who have donated tissue for scientific research and their families who have supported this. This study was supported by the Academy of Medical Sciences Springboard Award (SBF006\1038) (S.D., and E.E.P.), UKRI Future Leaders Fellowship (MR/V023861/1) (E.F.G., A.U., and S.D.), Parkinson’s UK Senior Fellowship (F1301) (H.M.), NZP UK Ltd (L.H., R.H., O.B., H.M.) and and DiMeN Doctoral Training Partnership award (MR/ N013840/1) (H.M., O.B. and R.H.). This study/research was also funded by the National Institute for Health and Care Research (NIHR) Sheffield Biomedical Research Centre (BRC). The views expressed are those of the authors and not necessarily those of the NIHR or the Department of Health and Social Care.

## Author contributions

E.E.P. conducted the majority of the cellular experiments, with assistance from F.C., D.P.R., L.H., A.S., E.S., and R.H. Kinetic modeling was performed by J.W. Histopathological analysis and tissue homogenization were carried out by E.F.G. and A.U. W.H.M. expressed and purified Bri2 BRICHOS, and M.O.C. performed mass spectrometry. J.R.H. assisted with tissue selection and provided postmortem samples. Data analysis, interpretation, and statistical evaluation were performed by E.E.P., J.W., E.F.G., A.U., and S.D. H.M. supervised the reprogramming, cell culture, and development of all cellular model lines. T.P.J.K., designed and supervised the kinetic modelling work. The overall project was supervised by S.D. with additional guidance from T.P.J.K., H.M., O.B., M.O.L., S.L., A.A., J.J., M.V., and J.R.H. The study was conceived and designed by E.E.P., and S.D. The first draft of the manuscript was written by E.E.P., and S.D. and all authors reviewed the manuscript, provided critical feedback, and approved the final version of this work.

## Reference

1. Poewe, W., Seppi, K., Tanner, C.M., Halliday, G.M., Brundin, P., Volkmann, J., Schrag, A.-E., and Lang, A.E. (2017). Parkinson disease. Nat. Rev. Dis. Prim. 3, 17013. 10.1038/nrdp.2017.13.

2. Bloem, B.R., Okun, M.S., and Klein, C. (2021). Parkinson’s disease. Lancet 397, 2284–2303. 10.1016/S0140-6736(21)00218-X.

3. Klein, C., and Westenberger, A. (2012). Genetics of Parkinson’s disease. Cold Spring Harb. Perspect. Med. 2, a008888. 10.1101/cshperspect.a008888.

4. Singleton, A.B., Farrer, M., Johnson, J., Singleton, A., Hague, S., Kachergus, J., Hulihan, M., Peuralinna, T., Dutra, A., Nussbaum, R., et al. (2003). α-Synuclein Locus Triplication Causes Parkinson’s Disease. Science (80-.). 302, 841. 10.1126/science.1090278.

5. Spillantini, M.G., Schmidt, M.L., Lee, V.M.-Y., Trojanowski, J.Q., Jakes, R., and Goedert, M. (1997). α-Synuclein in Lewy bodies. Nature 388, 839–840. 10.1038/42166.

6. Polymeropoulos, M.H., Lavedan, C., Leroy, E., Ide, S.E., Dehejia, A., Dutra, A., Pike, B., Root, H., Rubenstein, J., Boyer, R., et al. (1997). Mutation in the α-Synuclein Gene Identified in Families with Parkinson’s Disease. Science (80-.). 276, 2045–2047. 10.1126/science.276.5321.2045.

7. Sasaki, S., Shirata, A., Yamane, K., and Iwata, M. (2004). Parkin-positive autosomal recessive juvenile parkinsonism with α-synuclein-positive inclusions. Neurology 63, 678–682. 10.1212/01.WNL.0000134657.25904.0B.

8. Mori, H., Kondo, T., Yokochi, M., Matsumine, H., Nakagawa-Hattori, Y., Miyake, T., Suda, K., and Mizuno, Y. (1998). Pathologic and biochemical studies of juvenile parkinsonism linked to chromosome 6q. Neurology 51, 890–892. 10.1212/WNL.51.3.890.

9. Coukos, R., and Krainc, D. (2024). Key genes and convergent pathogenic mechanisms in Parkinson disease. Nat. Rev. Neurosci. 25, 393–413. 10.1038/s41583-024-00812-2.

10. Mahul-Mellier, A.-L., Burtscher, J., Maharjan, N., Weerens, L., Croisier, M., Kuttler, F., Leleu, M., Knott, G.W., and Lashuel, H.A. (2020). The process of Lewy body formation, rather than simply α-synuclein fibrillization, is one of the major drivers of neurodegeneration. Proc. Natl. Acad. Sci. 117, 4971–4982. 10.1073/pnas.1913904117.

11. Fares, M.B., Jagannath, S., and Lashuel, H.A. (2021). Reverse engineering Lewy bodies: how far have we come and how far can we go? Nat. Rev. Neurosci. 22, 111–131. 10.1038/s41583-020-00416-6.

12. Shahmoradian, S.H., Lewis, A.J., Genoud, C., Hench, J., Moors, T.E., Navarro, P.P., Castaño-Díez, D., Schweighauser, G., Graff-Meyer, A., Goldie, K.N., et al. (2019). Lewy pathology in Parkinson’s disease consists of crowded organelles and lipid membranes. Nat. Neurosci. 22, 1099–1109. 10.1038/s41593-019-0423-2.

13. Spillantini, M.G., Crowther, R.A., Jakes, R., Hasegawa, M., and Goedert, M. (1998). α-Synuclein in filamentous inclusions of Lewy bodies from Parkinson’s disease and dementia with Lewy bodies. Proc. Natl. Acad. Sci. 95, 6469–6473. 10.1073/pnas.95.11.6469.

14. Fujiwara, H., Hasegawa, M., Dohmae, N., Kawashima, A., Masliah, E., Goldberg, M.S., Shen, J., Takio, K., and Iwatsubo, T. (2002). α-Synuclein is phosphorylated in synucleinopathy lesions. Nat. Cell Biol. 4, 160–164. 10.1038/ncb748.

15. Anderson, J.P., Walker, D.E., Goldstein, J.M., de Laat, R., Banducci, K., Caccavello, R.J., Barbour, R., Huang, J., Kling, K., Lee, M., et al. (2006). Phosphorylation of Ser-129 Is the Dominant Pathological Modification of α-Synucleinin Familial and Sporadic Lewy Body Disease. J. Biol. Chem. 281, 29739–29752. 10.1074/jbc.M600933200.

16. Lee, K.-W., Chen, W., Junn, E., Im, J.-Y., Grosso, H., Sonsalla, P.K., Feng, X., Ray, N., Fernandez, J.R., Chao, Y., et al. (2011). Enhanced Phosphatase Activity Attenuates α-Synucleinopathy in a Mouse Model. J. Neurosci. 31, 6963 LP – 6971. 10.1523/JNEUROSCI.6513-10.2011.

17. Ramalingam, N., Jin, S.-X., Moors, T.E., Fonseca-Ornelas, L., Shimanaka, K., Lei, S., Cam, H.P., Watson, A.H., Brontesi, L., Ding, L., et al. (2023). Dynamic physiological α-synuclein S129 phosphorylation is driven by neuronal activity. npj Park. Dis. 9, 4. 10.1038/s41531-023-00444-w.

18. Parra-Rivas, L.A., Madhivanan, K., Aulston, B.D., Wang, L., Prakashchand, D.D., Boyer, N.P., Saia-Cereda, V.M., Branes-Guerrero, K., Pizzo, D.P., Bagchi, P., et al. (2023). Serine-129 phosphorylation of α-synuclein is an activity-dependent trigger for physiologic protein-protein interactions and synaptic function. Neuron 111, 4006–4023.e10. 10.1016/j.neuron.2023.11.020.

19. Braak, H., Tredici, K. Del, Rüb, U., de Vos, R.A.I., Jansen Steur, E.N.H., and Braak, E. (2003). Staging of brain pathology related to sporadic Parkinson’s disease. Neurobiol. Aging 24, 197–211. 10.1016/S0197-4580(02)00065-9.

20. Attems, J., Toledo, J.B., Walker, L., Gelpi, E., Gentleman, S., Halliday, G., Hortobagyi, T., Jellinger, K., Kovacs, G.G., Lee, E.B., et al. (2021). Neuropathological consensus criteria for the evaluation of Lewy pathology in post-mortem brains: a multi-centre study. Acta Neuropathol. 141, 159–172. 10.1007/s00401-020-02255-2.

21. Brundin, P., Li, J.-Y., Holton, J.L., Lindvall, O., and Revesz, T. (2008). Research in motion: the enigma of Parkinson’s disease pathology spread. Nat. Rev. Neurosci. 9, 741–745. 10.1038/nrn2477.

22. Prymaczok, N.C., De Francesco, P.N., Mazzetti, S., Humbert-Claude, M., Tenenbaum, L., Cappelletti, G., Masliah, E., Perello, M., Riek, R., and Gerez, J.A. (2024). Cell-to-cell transmitted alpha-synuclein recapitulates experimental Parkinson’s disease. npj Park. Dis. 10, 10. 10.1038/s41531-023-00618-6.

23. Luk, K.C., Kehm, V., Carroll, J., Zhang, B., O’Brien, P., Trojanowski, J.Q., and Lee, V.M.-Y. (2012). Pathological α-Synuclein Transmission Initiates Parkinson-like Neurodegeneration in Nontransgenic Mice. Science (80-.). 338, 949–953. 10.1126/science.1227157.

24. Hansen, C., Angot, E., Bergström, A.-L., Steiner, J.A., Pieri, L., Paul, G., Outeiro, T.F., Melki, R., Kallunki, P., Fog, K., et al. (2011). α-Synuclein propagates from mouse brain to grafted dopaminergic neurons and seeds aggregation in cultured human cells. J. Clin. Invest. 121, 715–725. 10.1172/JCI43366.

25. Gustafsson, G., Lööv, C., Persson, E., Lázaro, D.F., Takeda, S., Bergström, J., Erlandsson, A., Sehlin, D., Balaj, L., György, B., et al. (2018). Secretion and Uptake of α-Synuclein Via Extracellular Vesicles in Cultured Cells. Cell. Mol. Neurobiol. 38, 1539–1550. 10.1007/s10571-018-0622-5.

26. Siderowf, A., Concha-Marambio, L., Lafontant, D.-E., Farris, C.M., Ma, Y., Urenia, P.A., Nguyen, H., Alcalay, R.N., Chahine, L.M., Foroud, T., et al. (2023). Assessment of heterogeneity among participants in the Parkinson’s Progression Markers Initiative cohort using α-synuclein seed amplification: a cross-sectional study. Lancet Neurol. 22, 407–417. 10.1016/S1474-4422(23)00109-6.

27. Coughlin, D.G., Shifflett, B., Farris, C.M., Ma, Y., Galasko, D., Edland, S.D., Mollenhauer, B., Brumm, M.C., Poston, K.L., Marek, K., et al. (2025). α-Synuclein Seed Amplification Assay Amplification Parameters and the Risk of Progression in Prodromal Parkinson Disease. Neurology 104, e210279. 10.1212/WNL.0000000000210279.

28. Bernhardt, A.M., Longen, S., Trossbach, S. V, Rossi, M., Weckbecker, D., Schmidt, F., Jäck, A., Katzdobler, S., Fietzek, U.M., Weidinger, E., et al. (2025). A quantitative Lewy-fold-specific alpha-synuclein seed amplification assay as a progression marker for Parkinson’s disease. Acta Neuropathol. 149, 20. 10.1007/s00401-025-02853-y.

29. Gatto, N., Dos Santos Souza, C., Shaw, A.C., Bell, S.M., Myszczynska, M.A., Powers, S., Meyer, K., Castelli, L.M., Karyka, E., Mortiboys, H., et al. (2021). Directly converted astrocytes retain the ageing features of the donor fibroblasts and elucidate the astrocytic contribution to human CNS health and disease. Aging Cell 20, e13281. 10.1111/acel.13281.

30. Bell, S.M., Wareing, H., Capriglia, F., Hughes, R., Barnes, K., Hamshaw, A., Adair, L., Shaw, A., Olejnik, A., De, S., et al. (2025). Increasing hexokinase 1 expression improves mitochondrial and glycolytic functional deficits seen in sporadic Alzheimer’s disease astrocytes. Mol. Psychiatry 30, 1369–1382. 10.1038/s41380-024-02746-8.

31. Schwartzentruber, A., Boschian, C., Lopes, F.M., Myszczynska, M.A., New, E.J., Beyrath, J., Smeitink, J., Ferraiuolo, L., and Mortiboys, H. (2020). Oxidative switch drives mitophagy defects in dopaminergic parkin mutant patient neurons. Sci. Rep. 10, 15485. 10.1038/s41598-020-72345-4.

32. Carling, P.J., Mortiboys, H., Green, C., Mihaylov, S., Sandor, C., Schwartzentruber, A., Taylor, R., Wei, W., Hastings, C., Wong, S., et al. (2020). Deep phenotyping of peripheral tissue facilitates mechanistic disease stratification in sporadic Parkinson’s disease. Prog. Neurobiol. 187, 101772. 10.1016/j.pneurobio.2020.101772.

33. Choi, M.L., Chappard, A., Singh, B.P., Maclachlan, C., Rodrigues, M., Fedotova, E.I., Berezhnov, A. V, De, S., Peddie, C.J., and Athauda, D. (2022). Pathological structural conversion of α-synuclein at the mitochondria induces neuronal toxicity. Nat. Neurosci. 25, 1134–1148.

34. Henrich, M.T., Oertel, W.H., Surmeier, D.J., and Geibl, F.F. (2023). Mitochondrial dysfunction in Parkinson’s disease – a key disease hallmark with therapeutic potential. Mol. Neurodegener. 18, 83. 10.1186/s13024-023-00676-7.

35. Jain, A., Liu, R., Ramani, B., Arauz, E., Ishitsuka, Y., Ragunathan, K., Park, J., Chen, J., Xiang, Y.K., and Ha, T. (2011). Probing cellular protein complexes using single-molecule pull-down. Nature 473, 484–488. 10.1038/nature10016.

36. Fauvet, B., and Lashuel, H.A. (2016). Semisynthesis and Enzymatic Preparation of Post-translationally Modified α-Synuclein BT - Protein Amyloid Aggregation: Methods and Protocols. In, D. Eliezer, ed. (Springer New York), pp. 3–20. 10.1007/978-1-4939-2978-8_1.

37. Cremades, N., Cohen, S.I.A., Deas, E., Abramov, A.Y., Chen, A.Y., Orte, A., Sandal, M., Clarke, R.W., Dunne, P., Aprile, F.A., et al. (2012). Direct observation of the interconversion of normal and toxic forms of α-synuclein. Cell 149, 1048–1059. 10.1016/j.cell.2012.03.037.

38. De, S., and Klenerman, D. (2019). Imaging individual protein aggregates to follow aggregation and determine the role of aggregates in neurodegenerative disease. Biochim. Biophys. Acta - Proteins Proteomics 1867, 870–878. 10.1016/j.bbapap.2018.12.010.

39. Fusco, G., Chen, S.W., Williamson, P.T.F., Cascella, R., Perni, M., Jarvis, J.A., Cecchi, C., Vendruscolo, M., Chiti, F., Cremades, N., et al. (2017). Structural basis of membrane disruption and cellular toxicity by α-synuclein oligomers. Science (80-.). 358, 1440–1443.

40. Emin, D., Zhang, Y.P., Lobanova, E., Miller, A., Li, X., Xia, Z., Dakin, H., Sideris, D.I., Lam, J.Y.L., Ranasinghe, R.T., et al. (2022). Small soluble α-synuclein aggregates are the toxic species in Parkinson’s disease. Nat. Commun. 13, 5512. 10.1038/s41467-022-33252-6.

41. Nakamura, K., Nemani, V.M., Azarbal, F., Skibinski, G., Levy, J.M., Egami, K., Munishkina, L., Zhang, J., Gardner, B., Wakabayashi, J., et al. (2011). Direct Membrane Association Drives Mitochondrial Fission by the Parkinson Disease-associated Protein α-Synuclein. J. Biol. Chem. 286, 20710–20726. 10.1074/jbc.M110.213538.

42. Flagmeier, P., De, S., Wirthensohn, D.C., Lee, S.F., Vincke, C., Muyldermans, S., Knowles, T.P.J.J., Gandhi, S., Dobson, C.M., and Klenerman, D. (2017). Ultrasensitive measurement of Ca2+ influx into lipid vesicles induced by protein aggregates. Angew. Chemie Int. Ed. 56, 7750–7754. 10.1002/anie.201700966.

43. Rodrigues, M., Bhattacharjee, P., Brinkmalm, A., Do, D.T., Pearson, C.M., De, S., Ponjavic, A., Varela, J.A., Kulenkampff, K., Baudrexel, I., et al. (2022). Structure-specific amyloid precipitation in biofluids. Nat. Chem. 14, 1045–1053. 10.1038/s41557-022-00976-3.

44. Sanderson, J.B., De, S., Jiang, H., Rovere, M., Jin, M., Zaccagnini, L., Hays Watson, A., De Boni, L., Lagomarsino, V.N., Young-Pearse, T.L., et al. (2020). Analysis of α-synuclein species enriched from cerebral cortex of humans with sporadic dementia with Lewy bodies. Brain Commun. 2, fca0010. 10.1093/braincomms/fcaa010.

45. Meisl, G., Kirkegaard, J.B., Arosio, P., Michaels, T.C.T., Vendruscolo, M., Dobson, C.M., Linse, S., and Knowles, T.P.J. (2016). Molecular mechanisms of protein aggregation from global fitting of kinetic models. Nat. Protoc. 11, 252–272. 10.1038/nprot.2016.010.

46. Xu, C.K., Meisl, G., Andrzejewska, E.A., Krainer, G., Dear, A.J., Castellana-Cruz, M., Turi, S., Edu, I.A., Vivacqua, G., Jacquat, R.P.B., et al. (2024). α-Synuclein oligomers form by secondary nucleation. Nat. Commun. 15, 7083. 10.1038/s41467-024-50692-4.

47. Paradies, G., Paradies, V., De Benedictis, V., Ruggiero, F.M., and Petrosillo, G. (2014). Functional role of cardiolipin in mitochondrial bioenergetics. Biochim. Biophys. Acta - Bioenerg. 1837, 408–417. 10.1016/j.bbabio.2013.10.006.

48. Ghio, S., Camilleri, A., Caruana, M., Ruf, V.C., Schmidt, F., Leonov, A., Ryazanov, S., Griesinger, C., Cauchi, R.J., Kamp, F., et al. (2019). Cardiolipin Promotes Pore-Forming Activity of Alpha-Synuclein Oligomers in Mitochondrial Membranes. ACS Chem. Neurosci. 10, 3815–3829. 10.1021/acschemneuro.9b00320.

49. Lurette, O., Martín-Jiménez, R., Khan, M., Sheta, R., Jean, S., Schofield, M., Teixeira, M., Rodriguez-Aller, R., Perron, I., Oueslati, A., et al. (2023). Aggregation of alpha-synuclein disrupts mitochondrial metabolism and induce mitophagy via cardiolipin externalization. Cell Death Dis. 14, 729. 10.1038/s41419-023-06251-8.

50. Adam, L., Kumar, R., Arroyo-Garcia, L.E., Molenkamp, W.H., Nowak, J.S., Klute, H., Farzadfard, A., Alkenayeh, R., Nielsen, J., Biverstål, H., et al. (2024). Specific inhibition of α-synuclein oligomer generation and toxicity by the chaperone domain Bri2 BRICHOS. Protein Sci. 33, e5091. 10.1002/pro.5091.

51. Ghosh, D., Torres, F., Schneider, M.M., Ashkinadze, D., Kadavath, H., Fleischmann, Y., Mergenthal, S., Güntert, P., Krainer, G., Andrzejewska, E.A., et al. (2024). The inhibitory action of the chaperone BRICHOS against the α-Synuclein secondary nucleation pathway. Nat. Commun. 15, 10038. 10.1038/s41467-024-54212-2.

52. Flagmeier, P., De, S., Michaels, T.C.T., Yang, X., Dear, A.J., Emanuelsson, Cecilia Vendruscolo, Michele Linse, S., Klenerman, D., Knowles, T.P.J., and Dobson, C.M. (2020). Direct measurement of lipid membrane disruption connects kinetics and toxicity of Aβ42 aggregation. Nat. Struct. Mol. Biol. 10, 886–891.

53. Dear, A.J., Michaels, T.C.T., Meisl, G., Klenerman, D., Wu, S., Perrett, S., Linse, S., Dobson, C.M., and Knowles, T.P.J. (2020). Kinetic diversity of amyloid oligomers. Proc. Natl. Acad. Sci. 117, 12087–12094. 10.1073/pnas.1922267117.

54. Sang, J.C., Hidari, E., Meisl, G., Ranasinghe, R.T., Spillantini, M.G., and Klenerman, D. (2021). Super-resolution imaging reveals α-synuclein seeded aggregation in SH-SY5Y cells. Commun. Biol. 4, 613. 10.1038/s42003-021-02126-w.

55. Karampetsou, M., Ardah, M.T., Semitekolou, M., Polissidis, A., Samiotaki, M., Kalomoiri, M., Majbour, N., Xanthou, G., El-Agnaf, O.M.A., and Vekrellis, K. (2017). Phosphorylated exogenous alpha-synuclein fibrils exacerbate pathology and induce neuronal dysfunction in mice. Sci. Rep. 7, 16533. 10.1038/s41598-017-15813-8.

56. Marotta, N.P., Ara, J., Uemura, N., Lougee, M.G., Meymand, E.S., Zhang, B., Petersson, E.J., Trojanowski, J.Q., and Lee, V.M.-Y. (2021). Alpha-synuclein from patient Lewy bodies exhibits distinct pathological activity that can be propagated in vitro. Acta Neuropathol. Commun. 9, 188. 10.1186/s40478-021-01288-2.

57. Uemura, N., Marotta, N.P., Ara, J., Meymand, E.S., Zhang, B., Kameda, H., Koike, M., Luk, K.C., Trojanowski, J.Q., and Lee, V.M.-Y. (2023). α-Synuclein aggregates amplified from patient-derived Lewy bodies recapitulate Lewy body diseases in mice. Nat. Commun. 14, 6892. 10.1038/s41467-023-42705-5.

58. Ebrahimi-Fakhari, D., Cantuti-Castelvetri, I., Fan, Z., Rockenstein, E., Masliah, E., Hyman, B.T., McLean, P.J., and Unni, V.K. (2011). Distinct Roles In Vivofor the Ubiquitin–Proteasome System and the Autophagy–Lysosomal Pathway in the Degradation of α-Synuclein. J. Neurosci. 31, 14508 LP – 14520. 10.1523/JNEUROSCI.1560-11.2011.

59. Decressac, M., Mattsson, B., Weikop, P., Lundblad, M., Jakobsson, J., and Björklund, A. (2013). TFEB-mediated autophagy rescues midbrain dopamine neurons from α-synuclein toxicity. Proc. Natl. Acad. Sci. 110, E1817–E1826. 10.1073/pnas.1305623110.

60. Pantazopoulou, M., Brembati, V., Kanellidi, A., Bousset, L., Melki, R., and Stefanis, L. (2021). Distinct alpha-Synuclein species induced by seeding are selectively cleared by the Lysosome or the Proteasome in neuronally differentiated SH-SY5Y cells. J. Neurochem. 156, 880–896. 10.1111/jnc.15174.

61. Cuervo, A.M., Stefanis, L., Fredenburg, R., Lansbury, P.T., and Sulzer, D. (2004). Impaired Degradation of Mutant α-Synuclein by Chaperone-Mediated Autophagy. Science (80-.). 305, 1292–1295. 10.1126/science.1101738.

62. Winslow, A.R., Chen, C.-W., Corrochano, S., Acevedo-Arozena, A., Gordon, D.E., Peden, A.A., Lichtenberg, M., Menzies, F.M., Ravikumar, B., Imarisio, S., et al. (2010). α-Synuclein impairs macroautophagy: implications for Parkinson’s disease. J. Cell Biol. 190, 1023–1037. 10.1083/jcb.201003122.

63. Bayati, A., Banks, E., Han, C., Luo, W., Reintsch, W.E., Zorca, C.E., Shlaifer, I., Del Cid Pellitero, E., Vanderperre, B., McBride, H.M., et al. (2022). Rapid macropinocytic transfer of α-synuclein to lysosomes. Cell Rep. 40. 10.1016/j.celrep.2022.111102.

64. Ryan, T., Bamm, V. V, Stykel, M.G., Coackley, C.L., Humphries, K.M., Jamieson-Williams, R., Ambasudhan, R., Mosser, D.D., Lipton, S.A., Harauz, G., et al. (2018). Cardiolipin exposure on the outer mitochondrial membrane modulates α-synuclein. Nat. Commun. 9, 817. 10.1038/s41467-018-03241-9.

65. Bentivenga, G.M., Mammana, A., Baiardi, S., Rossi, M., Ticca, A., Magliocchetti, F., Mastrangelo, A., Poleggi, A., Ladogana, A., Capellari, S., et al. (2024). Performance of a seed amplification assay for misfolded alpha-synuclein in cerebrospinal fluid and brain tissue in relation to Lewy body disease stage and pathology burden. Acta Neuropathol. 147, 18. 10.1007/s00401-023-02663-0.

66. Sullivan, P.M., Mezdour, H., Aratani, Y., Knouff, C., Najib, J., Reddick, R.L., Quarfordt, S.H., and Maeda, N. (1997). Targeted Replacement of the Mouse Apolipoprotein E Gene with the Common Human APOE3 Allele Enhances Diet-induced Hypercholesterolemia and Atherosclerosis. J. Biol. Chem. 272, 17972–17980. 10.1074/jbc.272.29.17972.

67. Bayati, A., Ayoubi, R., Aguila, A., Zorca, C.E., Deyab, G., Han, C., Recinto, S.J., Nguyen-Renou, E., Rocha, C., Maussion, G., et al. (2024). Modeling Parkinson’s disease pathology in human dopaminergic neurons by sequential exposure to α-synuclein fibrils and proinflammatory cytokines. Nat. Neurosci. 27, 2401–2416. 10.1038/s41593-024-01775-4.

68. Uversky, V.N., Li, J., and Fink, A.L. (2001). Metal-triggered Structural Transformations, Aggregation, and Fibrillation of Human α-Synuclein: A POSSIBLE MOLECULAR LINK BETWEEN PARKINSON′S DISEASE AND HEAVY METAL EXPOSURE. J. Biol. Chem. 276, 44284–44296. 10.1074/jbc.M105343200.

69. Rusilowicz-Jones, E. V, Barone, F.G., Lopes, F.M., Stephen, E., Mortiboys, H., Urbé, S., and Clague, M.J. (2022). Benchmarking a highly selective USP30 inhibitor for enhancement of mitophagy and pexophagy. Life Sci. Alliance 5, e202101287. 10.26508/lsa.202101287.

70. Drews, A., De, S., Flagmeier, P., Wirthensohn, D.C., Chen, W.-H., Whiten, D.R., Rodrigues, M., Vincke, C., Muyldermans, S., Paterson, R.W., et al. (2017). Inhibiting the Ca2+ influx induced by human CSF. Cell Rep. 21, 3310–3316.

71. Dear, A.J., Thacker, D., Wennmalm, S., Ortigosa-Pascual, L., Andrzejewska, E.A., Meisl, G., Linse, S., and Knowles, T.P.J. (2024). Aβ Oligomer Dissociation Is Catalyzed by Fibril Surfaces. ACS Chem. Neurosci. 15, 2296–2307. 10.1021/acschemneuro.4c00127.

