## Supporting info file for "Secondary nucleation of α-Synuclein drives Mitochondria dysfunctions and Lewy body formation in Parkinson’s Disease"

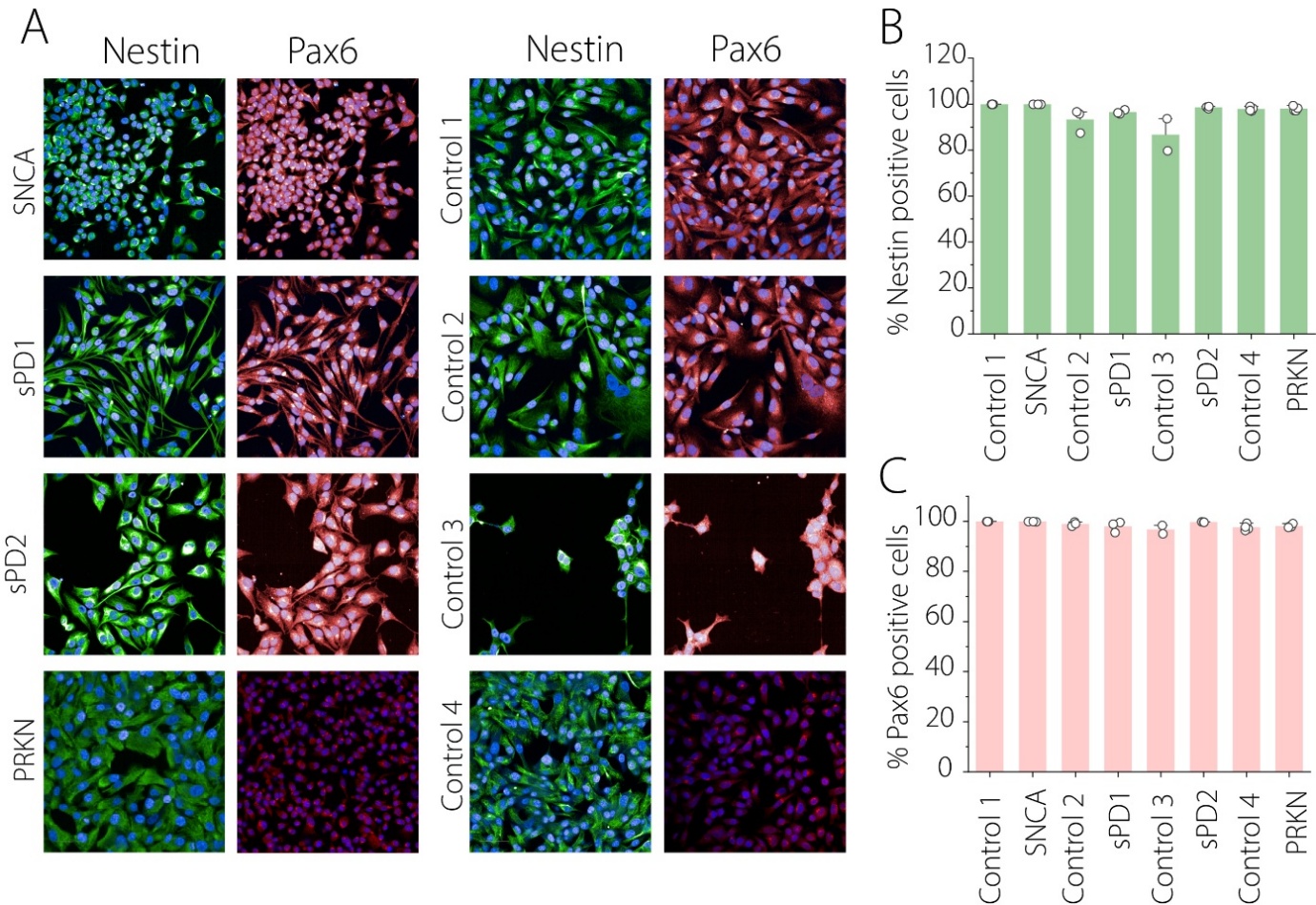

**Figure S1. (A)** Immunofluorescence staining of iNPCs for markers: Nestin and Pax6 in patient and NHC lines **(B-C)** Quantification of marker-positive cells as a percentage of total nuclei (Hoechst) with Nestin (B) and Pax6 (C)

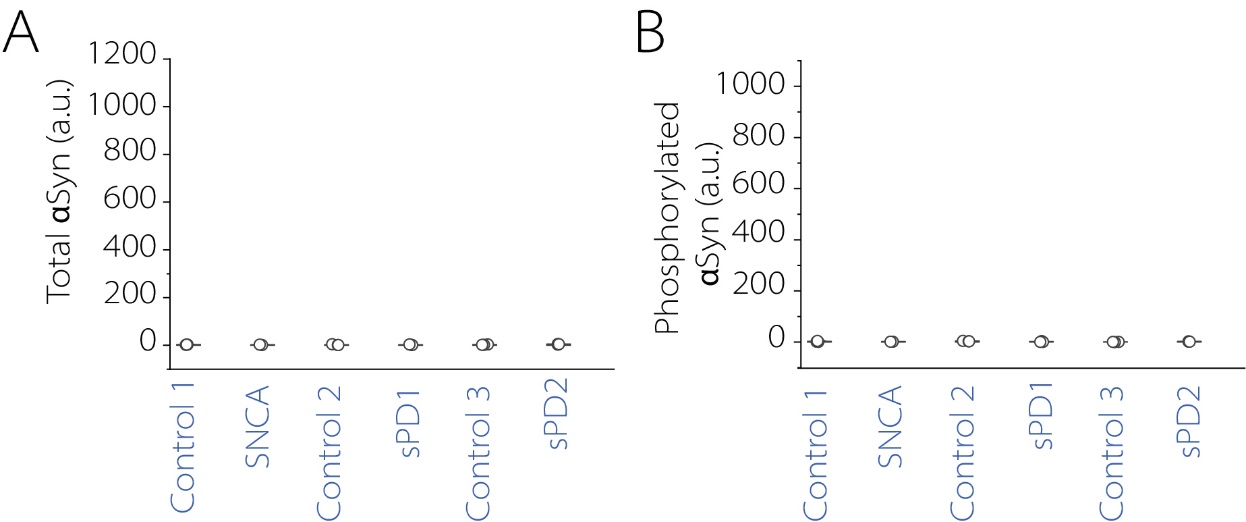

**Figure S2. (A -B)** Quantification of total αSyn(A), phosphorylated αSyn (pS129) (B), in fibroblast lysate across cell lines using an MSD assay.

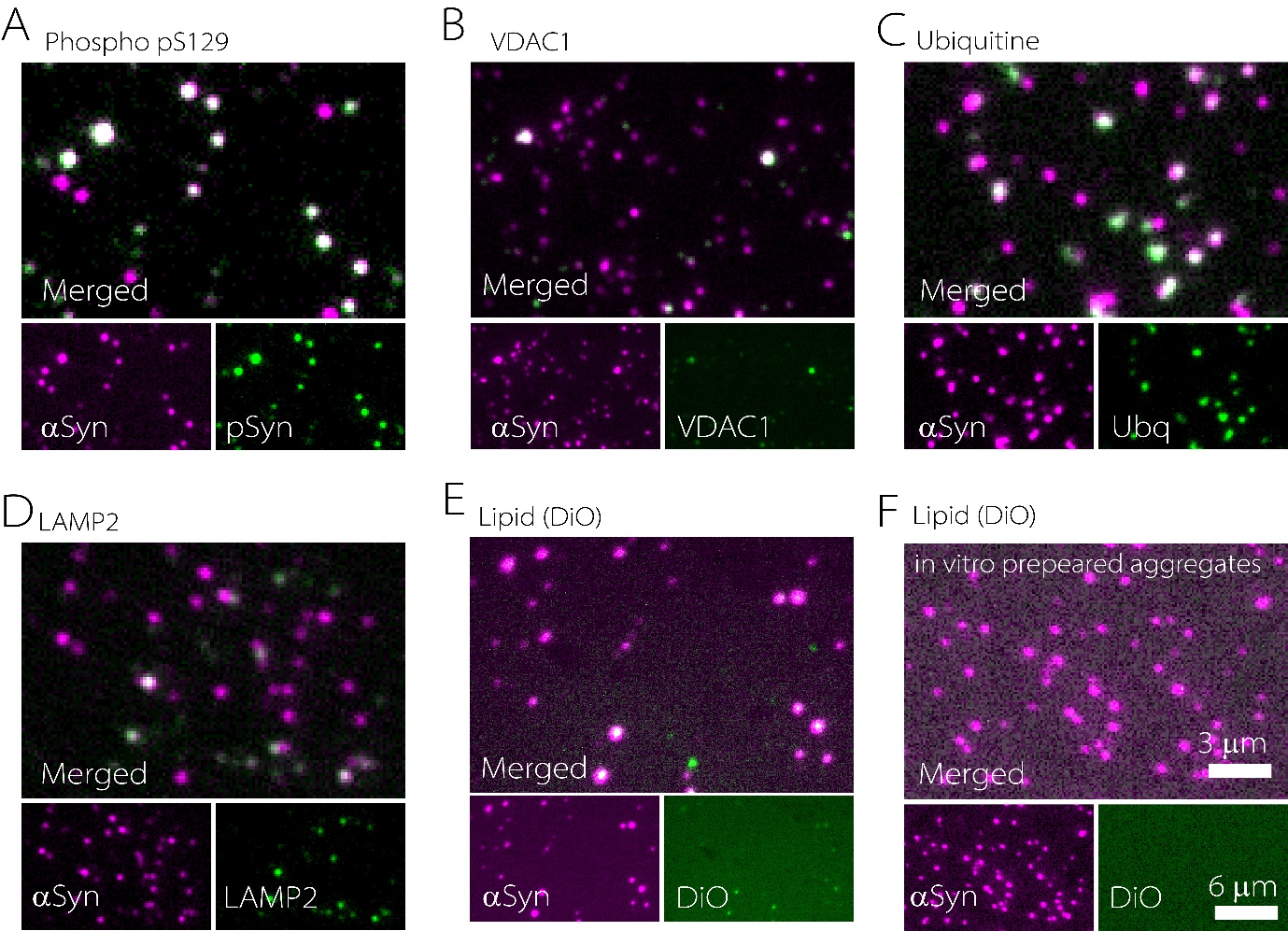

**Figure S3. (A-E)** Representative SiMPull images showing colocalization of αSyn aggregates with phosphorylated αSyn (pS129) (A), VDAC1 (B), Ubiquitin (C), LAMP2 (D), and Lipid (E) in iNL lysate. **(F)** Colocalization of recombinant αSyn aggregates with lipid marker DiO.

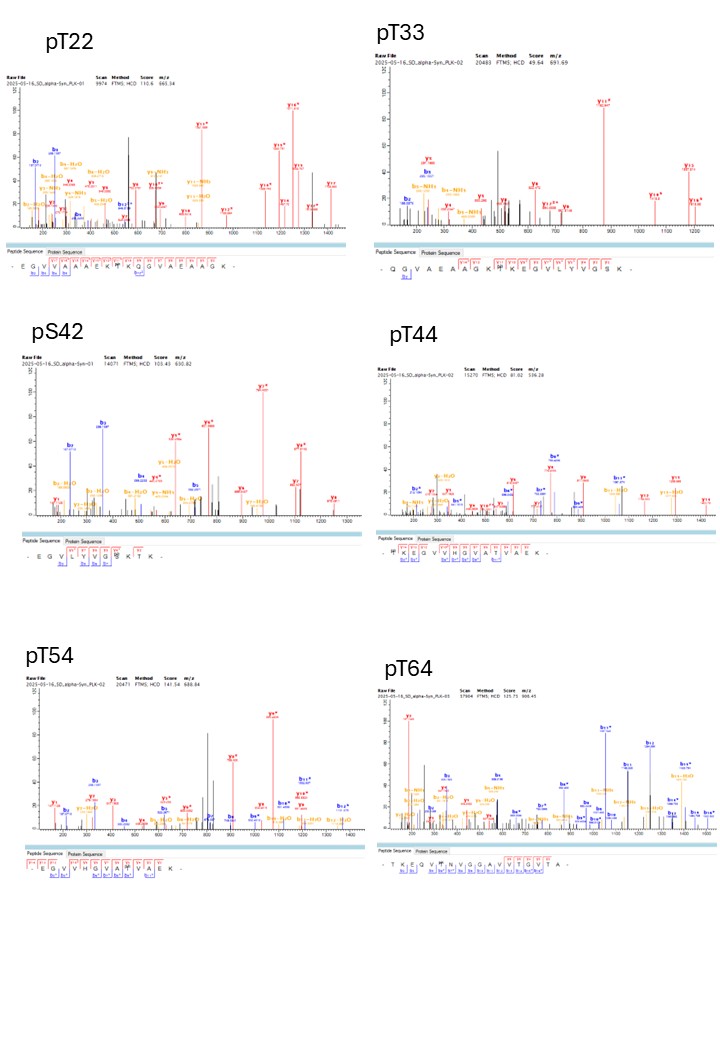

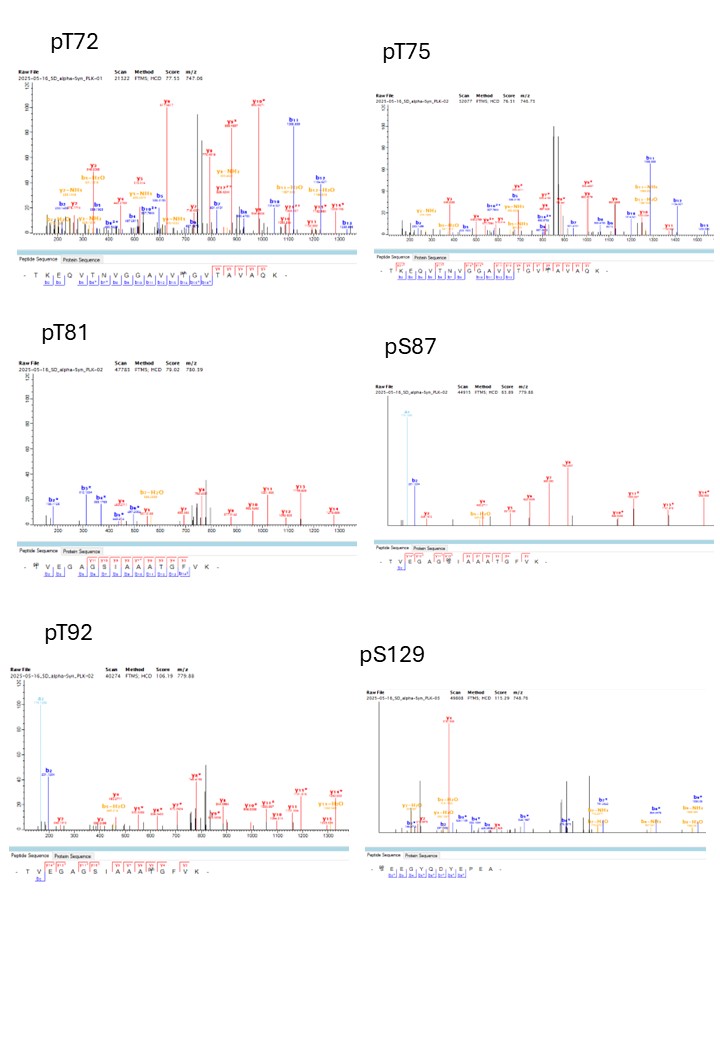

**Figure S4. Identification of phosphorylation sites using mass spectrometry.**

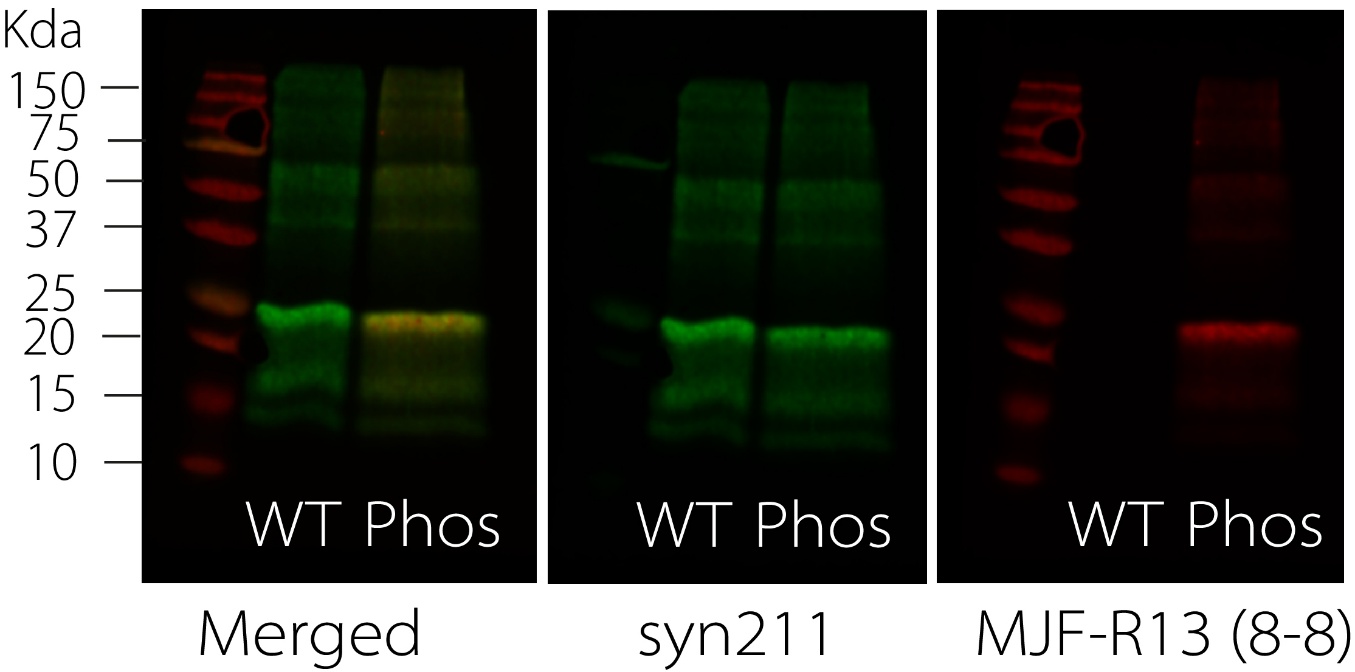

**Figure S5** Western blot confirming phosphorylation of AF488 labelled recombinant αSyn. WT αSyn was detected using Syn211 antibodies, and phosphorylated αSyn was detected using the phospho-specific MJF-R13 (8-8) antibody.

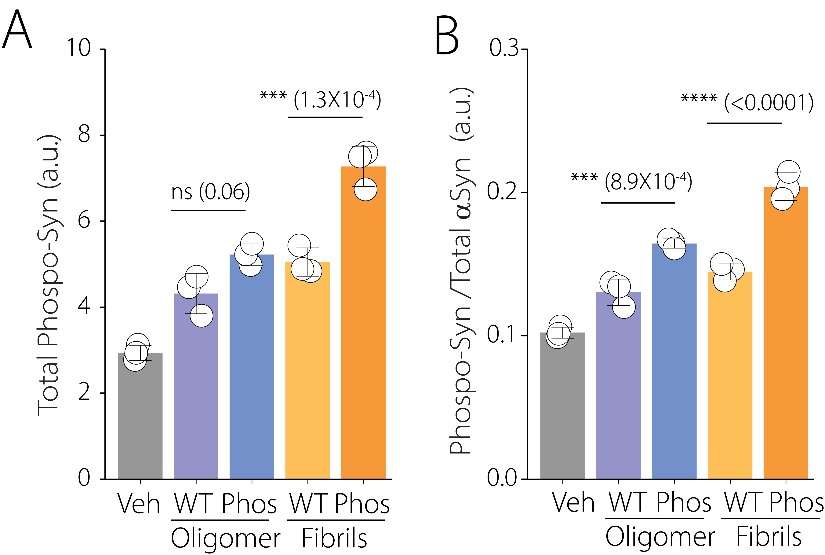

**Figure S6** Quantification of phosphorylated αSyn(A), and the ratio of phosphorylated αSyn and total αSyn (B) in lysate across cell lines using an MSD assay following different seed treatment.
